# A genetically-encoded fluorescent acetylcholine indicator

**DOI:** 10.1101/311126

**Authors:** Miao Jing, Peng Zhang, Guangfu Wang, Huoqing Jiang, Lukas Mesik, Jiesi Feng, Jianzhi Zeng, Shaohua Wang, Jess Looby, Nick A. Guagliardo, Linda W. Langma, Ju Lu, Yi Zuo, David A. Talmage, Lorna W. Role, Paula Q. Barrett, Li I. Zhang, Minmin Luo, Yan Song, J. Julius Zhu, Yulong Li

## Abstract

Acetylcholine (ACh) regulates a diverse array of physiological processes throughout the body, yet cholinergic transmission in the majority of tissues/organs remains poorly understood due primarily to the limitations of available ACh-monitoring techniques. We developed a family of <u>G</u>-protein-coupled receptor activation-based <u>ACh</u> sensors (GACh) with sensitivity, specificity, signal-to-noise ratio, kinetics and photostability suitable for monitoring ACh signals *in vitro* and *in vivo*. GACh sensors were validated with transfection, viral and/or transgenic expression in a dozen types of neuronal and non-neuronal cells prepared from several animal species. In all preparations, GACh sensors selectively responded to exogenous and/or endogenous ACh with robust fluorescence signals that were captured by epifluorescent, confocal and/or two-photon microscopy. Moreover, analysis of endogenous ACh release revealed firing pattern-dependent release and restricted volume transmission, resolving two long-standing questions about central cholinergic transmission. Thus, GACh sensors provide a user-friendly, broadly applicable toolbox for monitoring cholinergic transmission underlying diverse biological processes.

## INTRODUCTION

Acetylcholine (**ACh**), the first identified neurotransmitter, mediates cell-to-cell communication in the central and peripheral nervous systems, as well as non-neuronal systems^1–7^. Cholinergic projection neurons within the mammalian brain primarily originate in three major nuclear groups^1–5^, including the basal forebrain nuclei (**BF**), and the brainstem pedunculopontine and laterodorsal tegmental nuclei. Cholinergic neurons within these groups project widely throughout the cortical and subcortical domains, consistent with their involvement in complex brain functions, including attention, perception, associative learning and sleep/awake states. Additional smaller populations of cholinergic neurons scatter throughout other brain areas (e.g., the medial habenula (**MHb**) and the striatum), contributing to behaviors related to motion, motivation and stress^1, 3, 8^. Dysregulation of central cholinergic transmission is linked to a number of brain disorders, including Alzheimer’s disease, addiction, epilepsy, Parkinson’s disease, schizophrenia and depression. In the peripheral nervous and non-nervous systems, ACh is released by both neurons and non-neuronal cells to relay fast transmission at the neuromuscular junction and to regulate functions of a variety of other tissues and organs, including the heart, liver and pancreas^5–7^. Dysregulation of peripheral and non-neuronal cholinergic signals is associated with multiple pathological states, including cardiovascular disease, obesity, diabetes, immune deficiency and cancer.

Despite the significance of ACh signals in many, fundamental aspects of our physiology, cholinergic transmission in the majority of tissues and organs remain poorly understood, due primarily to the limitations of tools available for the direct measurement of ACh^1, 5, 9^. Microdialysis, an established method for monitoring extracellular ACh^10^, is less frequently used in current work because of its poor spatial and temporal resolution. Patch-clamp recordings of ACh-mediated currents have excellent sensitivity and temporal resolution, but the approach is limited by the number of cells that can be recorded and the prominent desensitization of cholinergic currents^3^. Similarly, ACh amperometry can achieve millisecond temporal resolution, yet this method is hampered by the technically challenging electrode enzymatic coating procedure that frequently ruins its stability and repeatability^11^. While TANGO assay has an unparalleled sensitivity, the time-consuming transcriptional and translational amplification processes make this approach inadequate for real-time ACh measurements^12^. Recent technical advances in fluorescence imaging have led to the development of FRET-based sensors, which allow rapid, optical detection of released ACh, although the low sensitivity limits their application *in vivo*^13, 14^. The latest cell-based fluorescent sensors (CNiFERs) allow live imaging of ACh dynamics in behaving animals^15, 16^. However, the dependence of CNiFERs on invasive cell transplantation limits broad use of this method. Nevertheless, the attractive real-time imaging features of both FRET-based sensors and CNiFERs inspired us to engineer more user-friendly and broadly applicable genetically-encoded ACh sensors^15, 17^.

Here, we report the development of a family of genetically-encoded G-protein-coupled receptor activation-based sensors for ACh (**GACh**). Our GACh sensors were initially constructed by coupling a circular permutated green fluorescent protein (**cpGFP**) into a muscarinic acetylcholine receptor (**MR**), with subsequent improvements via large-scale site-directed mutagenesis and screening. The sensitivity and utility of GACh sensors were validated in cultured cells, cultured cortical neurons, tissue slices prepared from multiple brain areas and peripheral organs, the olfactory system of living *Drosophila* and the visual cortex in awake behaving mice *in vivo*. Our data indicate that GACh sensors have the sensitivity (EC_50_ ≈ 1 μM), specificity (≈ MRs), signal-to-noise ratio (SNR ≈ 14), kinetics (*τ_on/off_* ≈ 200-800 ms) and photostability (≥ 1-4 hrs) suitable for precise and convenient real-time assays of ACh signals.

## RESULTS

### Development and optimization of GACh sensors

We initiated the development of a genetically-encoded optical sensor for ACh by inserting a conformationally sensitive cpGFP into all five subtypes of human muscarinic acetylcholine receptors (M_1-5_Rs) at their third intracellular loop (**ICL_3_**) (Fig. 1a). This site was chosen because ICL3 links the transmembrane helices 5 and 6 of muscarinic receptors and may undergo a large conformational change upon ACh binding^18^. To avoid creating a lengthy cpGFP-containing ICL_3_, that is likely to hinder the expression and surface trafficking of the receptors, and to minimize coupling of the receptors to downstream signaling, we replaced ICL_3_ of M_1-5_Rs with a shorter 24-amino acid ICL_3_ modified from an adrenergic β2R (**Fig. S1**). We then expressed all chimeras, M_1-5_R-β_2_R ICL_3_-cpGFP, in HEK293T cells to evaluate their response to ACh. Cells expressing the M_3_R chimera, M_3_R-β_2_R ICL_3_-cpGFP, which showed excellent membrane expression, had increased fluorescence responses (ΔF/F) (by ~30%) to bath application of ACh (Fig. 1b, red arrow head). In contrast, cells expressing the four other muscarinic receptor chimeras, all of which had poor membrane expression, exhibited no detectable ΔF/F upon ACh application (**Fig. S1b**). These results indicate that the M_3_R-derived sensor, that we called GACh1.0, can detect ACh.

**Figure 1:**
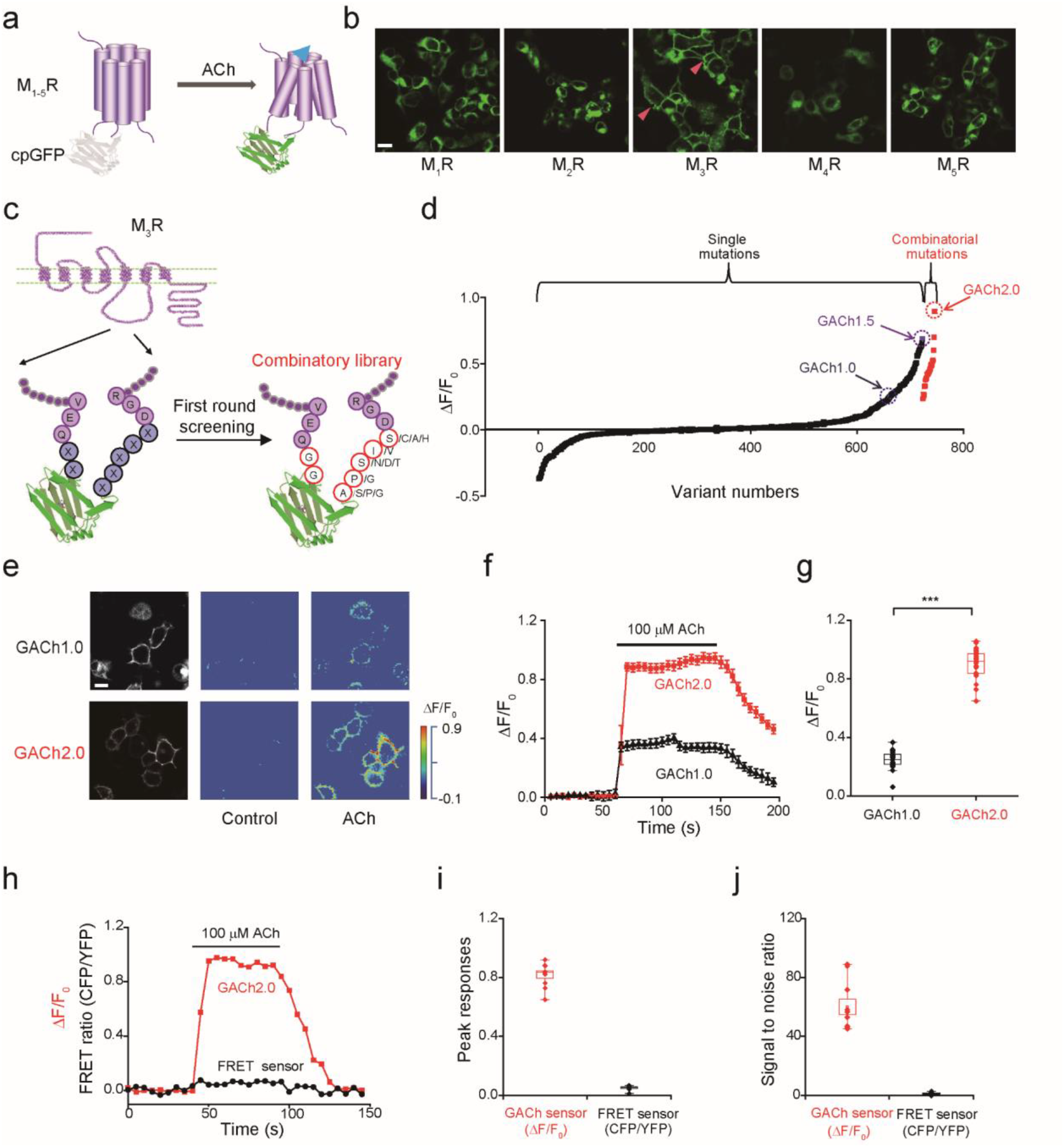
Development of GACh sensors. (**a**) Schematic drawing shows the principle of the GPCR Activation Based (GACh) sensor. (**b**) Membrane expression of the different MR-based candidate GACh sensors in HEK293T cells. Note the best membrane trafficking of M_3_R-based GACh1.0 sensor, marked by red arrow heads. (**c**) The top performers of the linker sequences of cpEGFP, including two amino acids in the N terminus and five amino acids in the C terminus. (**d**) Fluorescence responses of HEK293T cells expressing one of ~750 candidate GACh1.0 sensors containing either randomized point or combinatorial mutations to the bath application of 100 μM ACh. Note that combinatorial mutations yielded GACh2.0 with ΔF/F_0_ close to 100% and that each data point was the averaged responses of 2-10 cells. (**e**) Fluorescence responses of GACh1.0 and GACh2.0 expressing cells to the bath application of ACh. (**f-g**) ΔF/F_0_ of GACh1.0 and GACh2.0 expressing cells to ACh application (GACh1.0: 24.62 ± 1.51%, *n* = 19, GACh2.0: 90.12 ± 1.74%, *n* = 29, *U* = 551, *p* < 0.001). (**h**) Fluorescence response of a typical cell expressing either GACh2.0 or MR-based FRET sensor to the application of ACh (100 μM). (**i-j**) Averaged ΔF/F_0_ or ΔFRET ratio (GACh2.0: 94.0 ± 3.0%, *n* = 10; FRET: 6.6 ± 0.4%, *n* = 10, *U* = 100, *p* < 0.001) and SNR (GACh2.0: 60.0 ± 5.4, *n* = 10; FRET: 1.12 ± 0.21, *n* = 10, *U* =100, *p* < 0.001) of GACh2.0 and MR-based FRET sensor expressing cells to ACh application. *** indicates *p* < 0.001 (Mann-Whitney Rank Sum non-parametric test). All scale bars, 10 μm.

To improve GACh1.0, we used site-directed mutagenesis to create a library of 723 randomized point mutations at the two- and five-amino acid linkers in the N and C termini of cpGFP in GACh1.0 (Fig. 1c and **Fig. S2**, see **Methods** for details). We then individually expressed these variants in HEK293T cells and screened for candidates with larger ΔF/F to ACh application. The screening revealed that the best variant had ΔF/F increased by ~70% compared to GACh1.0, and we named the variant GACh1.5 (Fig. 1d). In addition, this screening revealed one or multiple single-point mutations for each of the seven linker residues (total 18 hits) that produced relatively larger ΔF/F responses to ACh application (Fig. 1c and **S2**). We initiated the second round of site-directed mutagenesis and screening, using combinations of the top hits for mutagenesis of the two-amino acid N terminus linker residues (i.e., GG) and 2-4 top hits for the five-amino acid C terminus linker residues (Fig. 1c and **S2**). After screening 23 combinatorial variants, we found that the variant, with the linker sequences of GG and APSVA, had the maximally enhanced ΔF/F response to ACh application, and we named this variant GACh2.0 (**Fig. S2**). Further analysis showed that GACh2.0 retained excellent expression and membrane trafficking properties (Fig. 1e), and had enhanced dynamic range (by 2.5-fold) compared to GACh1.0 (Fig. 1f, g; **Movie S1**). Compared with the FRET based ACh sensor^14^, GACh2.0 had ~20-fold larger peak signal responses and ~60-fold higher signal-to-noise ratio (**SNR**) to 100 μM ACh application (Fig. 1h-j and **Fig. S3**).

### Characterization of GACh sensors in cultured cells and neurons

To characterize the properties of GACh, we first performed high-speed line scan imaging of fluorescent signals in the membrane surface of GACh2.0 expressing HEK293T cells in response to agonists or antagonists delivered with a rapid local perfusion system (Fig. 2a-b). Local ACh perfusion elicited increases in fluorescence intensity in GACh2.0 expressing cells that exhibited a rapid time course, while local perfusion of tiotropium, a muscarinic antagonist^19^, decreased fluorescent signals in GACh2.0 expressing cells bathed in 100 μM ACh, albeit with a slower time course (Fig. 2b-c). The time-dependent changes in fluorescence intensity were fitted to single exponential functions, yielding average activation and inactivation time constants of 280±32 ms and 762±75 ms, respectively (Fig. 2b-c). These values were likely overestimated due to application delay caused by the perfusion system (~80 ms, **Fig. S4**).

**Figure 2:**
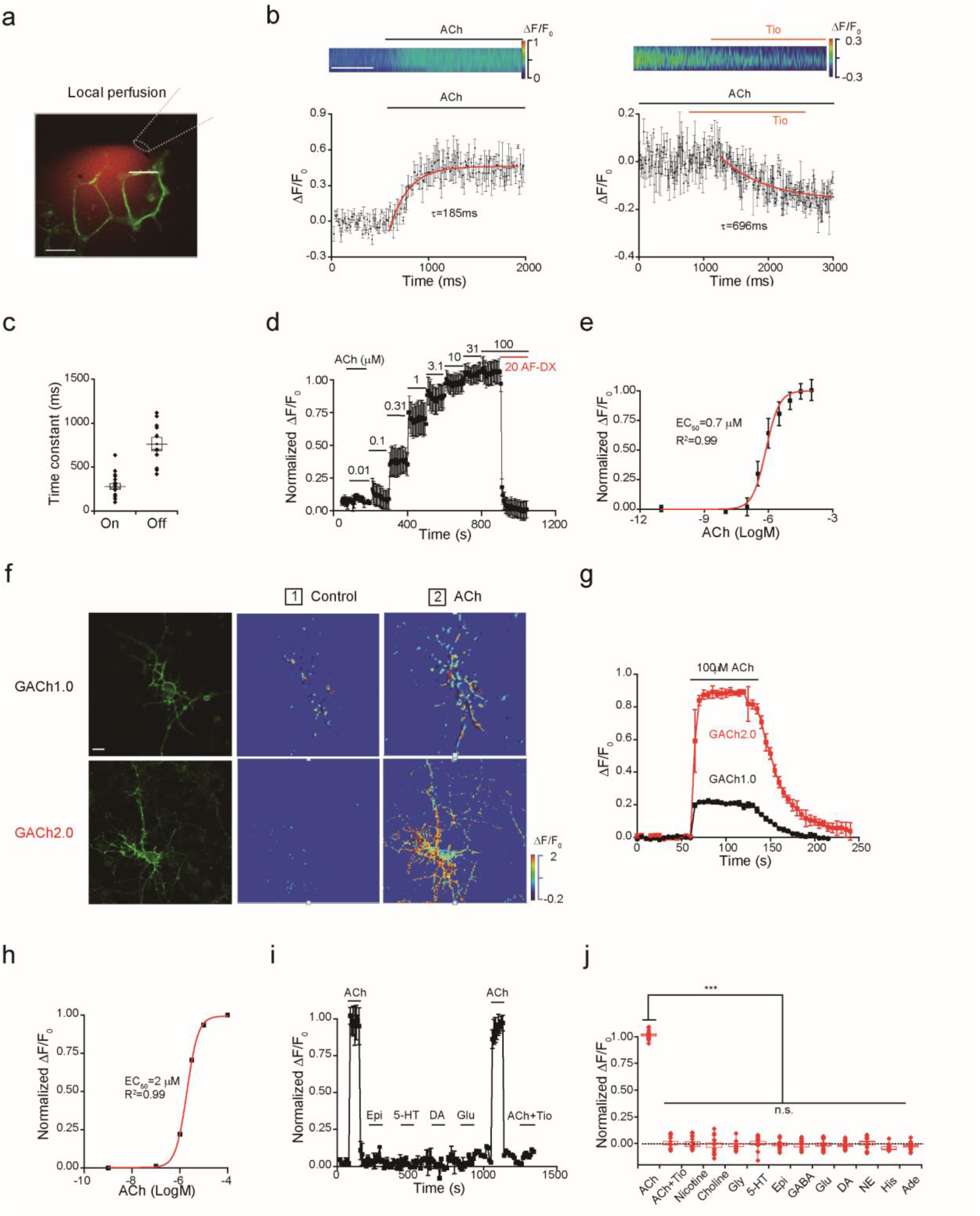
Characterization of GACh sensors in cultured HEK293T cells and neurons. (**a**) Illustration of a fast perfusion system, where a glass pipette filled with ACh and a red Rhodamine-6G dye was placed close to a GACh2.0 expressing cell. A white line indicated where the line scanning was performed. (**b**) Upper, scanning traces of fluorescence responses of GACh2.0 expressing cells to application of ACh and Tio. Lower plot shows fluorescence values of on and off responses of a GACh2.0 expressing cell to the application of ACh or Tio, averaged from 3 different ROIs on the scanning line. The original data was processed with 4 × 4 binning and plotted in (b). The white line indicates 0.5 s. (**c**) Averaged on (279.4 ± 32.6 ms, *n* = 18) and off (762.3 ± 74.9 ms, *n* = 11) time constants. Note that the time constants of GACh2.0 are likely overestimated due to the delay of perfusion system (~80 ms, **Fig. S4**). (**d**) Averaged responses (3 trials) of a GACh2.0 expressing HEK293T cell to ACh application. Note blockade of the responses by muscarinic antagonist AF-DX 384. (**e**) Dose-dependent response plot of GACh2.0 expressing HEK293T cells to ACh application yielded pEC_50_ = −6.12 ± 0.11 M, or EC_50_ = 0.78 ± 0.25 μM, *n* = 4). (**f**) Confocal GFP fluorescent and pseudocolor images of GACh1.0 and GACh2.0 expressing cultured cortical neurons in the normal bath solution and solution containing 100 μM ACh. (**g**) Time course of the fluorescence response of GACh1.0 and GACh2.0 expressing cultured neurons (averaged from 3 independent trials of one neuron). (**h**) Dose-dependent responses of GACh2.0 expressing cultured neurons (pEC_50_ = −5.70 ± 0.01 M or EC_50_ = 1.99 ±0.05 μM; *n* = 15). (**i**) Responses of GACh2.0 expressing neurons to application of ACh and other major neurotransmitters/modulators and ACh related compounds. (**j**) Values for normalized ΔF/F of GACh2.0 expressing neurons to application of 100 μM ACh with 2 μM tiotropium (Tio), 50 μM Nicotine, 100 μM Choline, 10 μM Glycine (Gly), 1 μM 5-HT, 10 μM Epinephrine (Epi), 10 μM GABA, 10 μM Glutamate (Glu), 20 μM Dopamine (DA), 200 μM Norepinephrine (NE), 1000 μM Histamine (His), 1 μM Adenosine (Ade) compared to application of ACh alone ( ACh: 90.99 ± 7.75%, *n* = 14; ACh+Tio: = 0.19 ± 1.53%, *n* = 14, *U* = 196, *p* < 0.001; Nicotine: 0.32 ± 1.47%, *n* = 15, *U* = 210, *p* < 0.001; Choline: −1.46 ± 2.31%, *n* = 15, *U* = 210, *p* < 0.001; Glycine: −1.36 ± 1.58%, *n* = 13, *U* = 182, *p* < 0.001; 5-HT: 0.96 ± 1.11%, *n* = 15, *U* = 210, *p* < 0.001; Epi: −0.77 ± 1.35%, *n* = 14, *U* = 196, *p* < 0.001; GABA: −2.01 ± 1.11%, *n* = 15, *U* = 210, *p* < 0.001; Glu: −0.49 ± 1.45%, *n* = 16, *U* = 224, *p* < 0.001; DA: −0.83 ± 1.20%, *n* = 15, *U* = 210, *p* < 0.001; NE: −0.42 ± 1.63%, *n* = 12, *U* = 168, *p* < 0.001; His: −4.54 ± 0.66%, *n* = 11, *U* = 154, *p* < 0.001; Ade: −2.23 ± 1.05%, *n* = 16, *U* = 224, *p* < 0.001; Mann-Whitney Rank Sum non-parametric tests with * indicating *p* < 0.05, ** indicating *p* < 0.01, and *** indicating *p* < 0.001, and n.s. indicating not significant).

To determine the sensor’s sensitivity to ACh, we measured the fluorescence intensity of GACh2.0 expressing HEK293T cells bathed in solutions containing varied ACh concentrations (Fig. 2d). Increasing ACh from 10 nM to 100 μM progressively increased the fluorescence intensity in GACh2.0 expressing cells. The concentration-response relationship was fitted to a Boltzmann equation with EC_50_ of ~0.7 μM (Fig. 2e), a value that is comparable to wild type M_3_Rs^20^. The ACh-induced fluorescent signals were completely blocked by co-application of 20 μM AF-DX 384, another muscarinic antagonist^21^, further indicating specificity.

Activation of muscarinic receptors stimulates intracellular arrestin and Ca^2+^ signaling^22, 23^. We compared the activation of GFP-tagged wild type M_3_Rs (GFP-M_3_Rs) and GACh2.0 expressed in HEK293T cells. Control GFP-M_3_R expressing cells had a significant reduction (~60%) in membrane GFP fluorescence intensity within 2 hours of ACh application, which was prevented by co-application of tiotropium (**Fig. S5a**), consistent with a muscarinic receptor-stimulated arrestin-dependent internalization^22^. By contrast, surface expression of GACh2.0 remained unchanged in the presence of 1 mM ACh for as long as 4 hours (**Fig. S5a**), indicating little internalization. In addition, using an arrestin TANGO assay^12^, we found that bath application of varied concentrations of ACh induced much smaller (~1000-fold less) reporter signals in GACh2.0 expressing cells compared to control GFP-M_3_R expressing cells (**Fig. S5b**). Together, these results suggest negligible coupling between GACh2.0 and downstream arrestin signaling events. Moreover, we examined the ACh-induced concentration-dependent Ca^2+^ responses in both GACh2.0 and GFP-M_3_R expressing cells (**Fig. S6a-c**). The Ca^2+^ responses in GACh2.0 expressing cells were ~7-fold smaller than those in GFP-M_3_R expressing cells (**Fig. S5c-e**), suggesting reduced coupling of GACh to downstream Gq protein-dependent signaling^23^. Collectively, these results indicate that GACh2.0 can sensitively detect ACh but minimally transduces intracellular signals.

To characterize GACh’s properties in neurons, we expressed GACh1.0 and GACh2.0 in cultured cortical neurons (Fig. 2f-j). Approximately 48 hours after transfection, GACh1.0 and GACh2.0 were distributed to throughout expressing neurons, with the majority of sensors delivered to the neurites (Fig. 2f). Bath application of 100 μM ACh enhanced the fluorescence intensity in cortical neurons expressing GACh1.0 and GACh2.0 by ~30% and ~90%, respectively (Fig. 2g; **Movie S2**), validating the functionality of the sensors in neurons. Varying ACh concentration in the bath solution from 1 nM to 100 μM progressively increased the fluorescence intensity in GACh2.0 expressing neurons, revealing a detectable signal at ~500 nM with an EC_50_ of ~2 μM (Fig. 2h).

To test the selectivity of GACh2.0 in neurons, we compared responses of GACh2.0 expressing neurons to bath application of ACh, other neurotransmitters and ACh-related bio-compounds. Bath application of ACh induced a pronounced increase in fluorescence intensity in GACh2.0 expressing neurons that was completely blocked by including additional 2 μM tiotropium in the bath solution. In contrast, bath application of nicotine, choline, glycine, serotonin, epinephrine, GABA, glutamate, dopamine, norepinephrine, histamine, and adenosine did not induce detectable change in fluorescence in GACh2.0 expressing neurons (Fig. 2i,j). Finally, we noted no alteration in membrane fluorescence intensity in GACh2.0 expressing neurons during a 30-minute bath application of 100 μM ACh (**Fig. S6a-c**), consistent with the minimal arrestin-dependent internalization of GACh2.0 in HEK293T cells. Together, these results support the utility of GACh as specific ACh sensors in neurons.

### Validation of GACh sensors in cultured brain slices

To further test the applicability of GACh sensors, we expressed the sensors in CA1 pyramidal neurons in cultured mouse hippocampal slices. We first used a lentiviral expression system to express GACh sensors in CA1 neurons for 7-14 days to provide sufficient time for GACh expression and trafficking to distal dendrites (**Fig. S7**). Two-photon imaging showed that GACh1.0, GACh1.5 or GACh2.0 were expressed throughout CA1 pyramidal neurons, with significant expression at plasma membrane of somata, dendrites and spines, sites of excitatory synapses (**Fig. S7**). To examine whether GACh sensors can detect ACh in the cultured slice preparation, we chose a Sindbis viral expression system that allowed a more rapid (~18 hours) and robust expression of GACh sensors in CA1 neurons (**Fig. S8a**; see the methods). Using an epifluorescent microscope, we captured fluorescence responses in GACh expressing neurons. A brief 500-ms puff application of ACh or a muscarinic agonist, oxotremorine^20^, evoked fluorescence responses in CA1 neurons expressing GACh1.0, GACh1.5 or GACh2.0, whereas puff application of a nicotinic agonist, nicotine, or control bath solution ACSF, induced no responses in the same neurons (**Fig. S8b-d**; **Movie S3**), indicating that GACh sensors selectively responded to ACh or muscarinic agonists.

Because GACh2.0 produced the largest ΔF/F to ACh in the above tests, we focused our investigation mainly on this sensor in the following experiments. We found that multiple puffs induced equivalent fluorescence responses in GACh2.0 expressing CA1 neurons (**Fig. S9**), suggesting photostability of GACh2.0. Bath application of 1 μM atropine, a muscarinic antagonist^21^, but not 2,2,6,6-tetramethylpiperidin-4-yl heptanoate, a nicotinic antagonist^24^, completely blocked ACh-induced fluorescence responses in GACh2.0 expressing CA1 neurons (**Fig. S9**), again indicating a specific muscarinic effect. Importantly, we found that the resting membrane potential, input resistance, membrane time constant and average spiking frequency of expressing GACh2.0 CA1 neurons were not different from nearby control non-expressing neurons (**Fig. S10a-d**), suggesting no effect of GACh2.0 expression on basic membrane properties. Similarly, AMPA, NMDA or GABAergic responses, or paired pulse facilitation of AMPA responses in GACh2.0 expressing CA1 neurons remained unchanged (**Fig. S10e-h**), indicating no alteration of synaptic transmission by GACh2.0 expression. Collectively, these results are consistent with the finding that GACh2.0 is a selective, photostable ACh sensor.

To compare the approaches of GACh2.0 imaging and patch-clamp recordings, we simultaneously made dual whole-cell recordings and fluorescence imaging from a pair of neighboring GACh2.0 expressing and control non-expressing CA3 pyramidal neurons in cultured mouse hippocampal slices (Fig. 3a); CA3 pyramidal neurons were chosen because they express relatively more endogenous muscarinic and nicotinic receptors and respond robustly to cholinergic activation^25^. A brief 500-ms ACh puff evoked a brief, large inward current followed by a prolonged, small inward current in both GACh2.0 expressing and control non-expressing CA3 neurons, presumably representing the responses of endogenous nicotinic and muscarinic receptors, respectively^3^ (Fig. 3b-d). A large concurrent fluorescent signal was observed only in GACh2.0 expressing neurons, but not in control non-expressing CA3 neurons (Fig. 3b-d). The latencies of cholinergic current responses and fluorescence signals were the same in GACh2.0 expressing neurons (Fig. 3b,e), indicating that GACh2.0 detected ACh as fast as endogenous cholinergic receptors. The SNR of GACh2.0 fluorescence responses (~14) was smaller than that of the fast nicotinic-like cholinergic current responses (~35), but larger than that of the slow muscarinic-like cholinergic current responses (~8) (Fig. 3b,f), indicating that GACh2.0 had a relatively comparable resolving power compared to the patch-clamp recordings in monitoring cholinergic signals. Notably, the second ACh puff evoked the same fluorescence responses, but smaller cholinergic currents (reduced by ~40%) in GACh2.0 expressing CA3 neurons compared to the first ACh puff (Fig. 3b,g,h), due presumably to desensitization of endogenous receptors^3^. There was no difference in the amplitude, latency or SNR of cholinergic currents in GACh2.0 expressing and control non-expressing CA3 neurons (Fig. 3b-f), confirming that expression of GACh2.0 had little non-specific effect on CA3 neurons. These results suggest that GACh2.0 is suitable for repeatedly and faithfully monitoring ACh signals.

**Figure 3:**
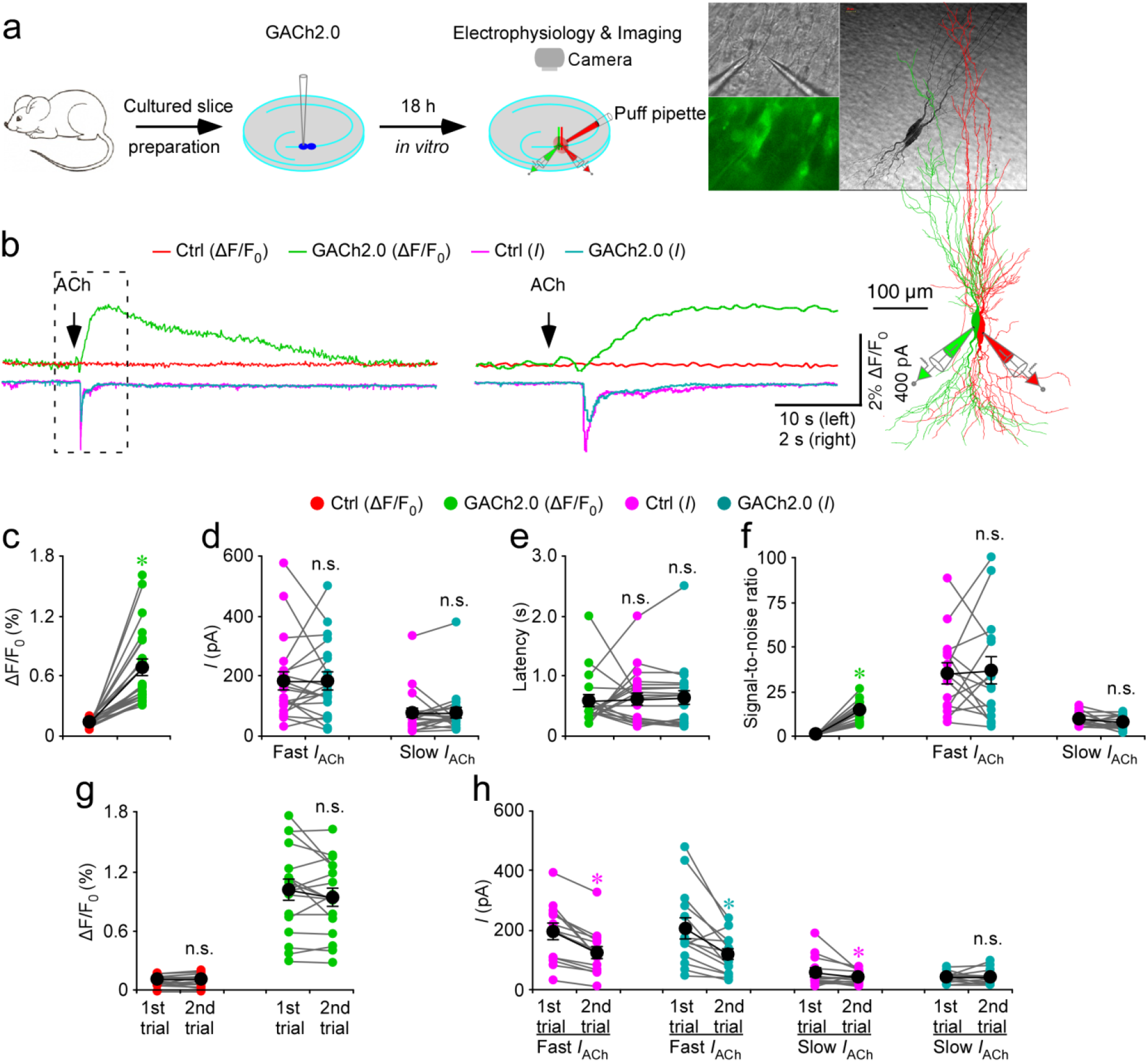
GACh2.0 detects rapid ACh application in brain slices. (**a**) Schematic drawing outlines the design of simultaneous imaging and electrophysiological recording experiments in mouse cultured hippocampal slice preparation. Left inserts show transmitted light (top), fluorescence microscopic (bottom) images of a pair of simultaneously recorded GACh2.0 expressing and neighboring control non-expressing CA3 neurons. Right inserts show the biocytin-filled and reconstructed GACh2.0 expressing and non-expressing CA3 neurons. (**b**) Left, simultaneous fluorescence and current responses of the pair of GACh2.0 expressing and neighboring control non-expressing CA3 neurons to a brief puff (500 ms) application of 100 mM acetylcholine (**ACh**). Right, the responses in the left rectangle box are shown again in an expanded time scale. Note the same latency of fluorescence and current responses. (**c**) Values for the cholinergic fluorescence responses of GACh2.0 expressing CA3 neurons compared to non-expressing neurons (GACh2.0: 0.68 ± 0.08%; Ctrl: 0.14 ± 0.01%; *Z* = 4.015; *p* < 0.001; *n* = 21 from 9 animals). (**d**) Values for the amplitudes of fast cholinergic current responses (GACh2.0: 180.9 ± 30.8 pA; Ctrl: 181.2 ± 28.4 pA; *Z* = −0.037; *p* = 0.97; *n* = 21 from 9 animals) and slow cholinergic current responses (GACh2.0: 76.2 ± 15.9 pA; Ctrl: 76.8 ± 17.0 pA; Z = 0.896; *p* = 0.37; *n* = 21 from 9 animals) in GACh2.0 expressing CA3 neurons compared to non-expressing neurons. (**e**) Values for the latencies of cholinergic current responses in non-expressing CA3 neurons (Ctrl: 611 ± 10 ms; *Z* = 0.523; *p* = 0.60) and GACh2.0 expressing (GACh2.0: 622 ± 12 ms; *Z* = 0.485; *p* = 0.62) compared to those of fluorescence responses of GACh2.0 expressing neurons (GACh2.0: 580 ± 9 ms; *n* = 21 from 9 animals). (**f**) Values for the signal-to-noise ratio (SNR) of cholinergic fluorescence responses of GACh2.0 expressing CA3 neurons compared to non-expressing neurons (GACh2.0: 14.0 ± 1.5; Ctrl: 1.0 ± 0.1; *Z* = 3.408; *p* < 0.005; *n* = 15 from 6 animals), and that of fast (GACh2.0: 36.2 ± 7.7; Ctrl: 34.6 ± 5.7; *Z* = 0.170; *p* = 0.86; *n* = 15 from 6 animals) and slow (GACh2.0: 7.5 ± 1.0; Ctrl: 9.0 ± 1.0; *Z* = −0.852; *p* = 0.39; *n* = 15 from 6 animals) cholinergic current responses of GACh2.0 expressing CA3 neurons compared to non-expressing neurons. Note that SNR of cholinergic fluorescence responses of GACh2.0 expressing CA3 neurons is smaller than fast (GACh2.0: *Z* = 2.242; *p* < 0.05; Ctrl: *Z* = 3.124; *p* < 0.005), but larger than slow (GACh2.0: *Z* = −2.840; *p* < 0.01; Ctrl: *Z* = −2.669; *p* < 0.01) cholinergic current responses of GACh2.0 expressing CA3 neurons compared to non-expressing neurons. (**g**) Values for the two fluorescence responses of non-expressing (1^st^: 0.11 ± 0.01%; 2^nd^: 0.11 ± 0.01%; *Z* = −0.142; *p* = 0.89; *n* = 17 from 9 animals) and GACh2.0 expressing (1^st^: 1.01 ± 0.11%; 2^nd^: 0.94 ± 0.09%; *Z* = −1.138; *p* = 0.26; *n* = 17 from 9 animals) CA3 neurons. (**h**) Values for the two fast cholinergic current responses in non-expressing (1^st^: 190.9 ± 26.1 pA; 2^nd^: 124.1 ± 20.4 pA; *Z* = −3.296; *p* < 0.005; *n* = 17 from 9 animals) and GACh2.0 expressing (1^st^: 203.8 ± 34.9 pA; 2^nd^: 119.3 ± 18.6 pA; *Z* = −2.856; *p* < 0.005; *n* = 17 from 9 animals) CA3 neurons, and values for the two slow cholinergic current responses of non-expressing (1^st^: 56.4 ± 13.4 pA; 2^nd^: 39.0 ± 5.7 pA; *Z* = −2.166; *p* < 0.05; *n* = 17 from 9 animals) and GACh2.0 expressing (1^st^: 41.6 ± 4.5 pA; 2^nd^: 41.7 ± 6.8 pA; *Z* = 0.940; *p* = 0.93; *n* = 17 from 9 animals) CA3 neurons. Large black dots indicate average responses and asterisks indicate *p* < 0.05 (Wilcoxon tests).

### Applications of GACh sensors in acute brain slices

To examine whether GACh2.0 detects ACh when expressed in the other brain areas and other animal species, we made *in vivo* Sindbis viral expression of GACh2.0 in a number of preparations, including layer 2 (**L2**) stellate neurons and L1 interneurons in the medial entorhinal cortex (**MEC**), L5 pyramidal neurons in the barrel cortex of mice, GABAergic thalamic reticular neurons and glutamatergic thalamocortical neurons in the ventral basal nucleus of rats. Approximately 18 hours after *in vivo* expression, we prepared acute brain slices containing these brain regions and measured ΔF/F to a brief puff application of ACh (**Fig. S11a**). Puff applications of ACh and oxotremorine, but not of nicotine or bath solution ACSF, evoked fluorescence increases in GACh2.0 expressing entorhinal L2 stellate neurons and L1 interneurons, barrel cortical L5 pyramidal neurons, thalamic reticular neurons and thalamocortical neurons (**Fig. S11b-c**). Collectively, these results suggest that GACh2.0 can selectively detect ACh in various brain areas of both mice and rats.

To test whether GACh2.0 can report endogenously released ACh, we used the same ***in vivo*** viral expression and ***ex vivo*** acute mouse slice preparation to measure ΔF/F responses of GACh2.0 expressing entorhinal L2 stellate neurons to electrical stimulation of MEC L1 (Fig. 4a), a layer that is densely innervated by cholinergic fibers originating from BF^26, 27^. Twenty pulses at 2-Hz evoked robust fluorescence responses in GACh2.0 expressing neurons (Fig. 4b,c; **Movie S4**). Repeated electric stimuli delivered every 8 minutes induced the same ΔF/F responses in GACh2.0 expressing neurons (Fig. 4d-e), indicating that GACh2.0 is suitable for monitoring endogenously released ACh signals over relatively long durations. Systematically varying the stimulation frequency revealed that low frequency stimuli (0.5–2 Hz) evoked large, plateau-like fluorescence responses, intermediate frequency stimuli (4–12 Hz) elicited more rapidly rising but briefer fluorescence responses, while high frequency stimuli (≥32 Hz) induced little fluorescence responses in GACh2.0 expressing neurons (Fig. 4f-h). Previous studies have shown that BF cholinergic neurons in behaving animals prefer low frequency (~0.5–2 Hz) tonic firing and theta rhythmic (~4—12 Hz) phasic firing^28, 29^. Our results suggest that these two preferred firing patterns generate distinct sustained and transient ACh release events, which are observed when animals perform different behavioral tasks^9^. Further altering the number of pulses delivered in 2-Hz electric stimuli showed that single electric pulses elicited detectable ΔF/F responses, whereas multiple pulses induced enhanced ΔF/F responses in GACh2.0 expressing neurons (Fig. 4i-k), suggesting that the amount of released ACh may scale the number of presynaptic action potentials.

**Figure 4:**
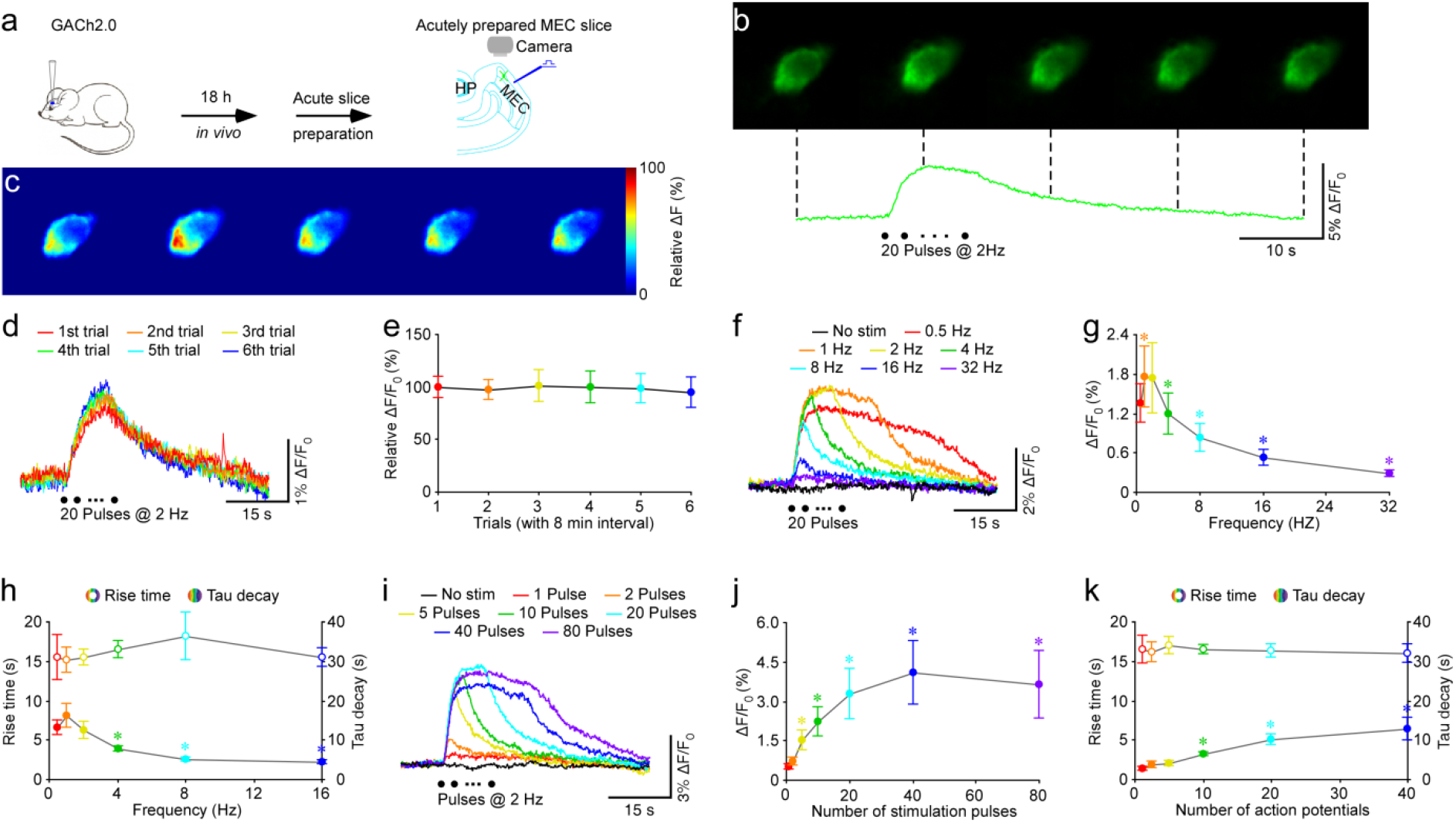
GACh2.0 reveals firing pattern-dependent ACh release modes in MEC. (**a**) Schematic drawing outlines the design of stimulation-imaging experiments in mouse MEC preparation. (**b**) Snapshots of fluorescence responses of a GACh2.0 expressing stellate cell to local electric stimuli. (**c**) Relative fluorescence responses of the GACh2.0 expressing stellate cell to local electric stimuli shown in a heat map format. (**d**) Fluorescence responses of a GACh2.0 expressing MEC stellate neuron to repetitive layer 1 electric stimulation every 8 minutes. (**e**) Values for the subsequent fluorescence responses of GACh2.0 MEC stellate neurons to the multiple layer 1 electric stimulation at time interval of 2 min (2^nd^: 1.58 ± 0.15%, *Z* = −0.534; *p* = 0.59; 3^rd^: 1.65 ± 0.25%, Z = −0.178; *p* = 0.86; 4^th^: 1.62 ± 0.25%, Z = 0.222; *p* = 0.82; 5^th^: 1.61 ± 0.22%, Z = 0.051; *p* = 0.96; 6^th^: 1.55 ± 0.23%, *Z* = −0.800; *p* = 0.42; *n* = 11 from 7 animals) compared to the first fluorescence response (1^st^: 1.63 ± 0.16%). (**f**) Fluorescence responses of a GACh2.0 expressing MEC stellate neuron to electric stimuli consisting of a train of 20 pulses at varied different frequency. (**g**) Values for the peak fluorescence responses of GACh2.0 expressing MEC stellate neurons to electric stimulations consisting of a train of 20 pulses at higher frequency (1 Hz: 1.75 ± 0.47%, *Z* = 2.606; *p* < 0.01; 2 Hz: 1.74 ± 0.53%, Z = 1.726; *p* = 0.08; 4 Hz: 1.19 ± 0.44%, Z = −1.746; *p* = 0.14; 8 Hz: 0.82 ± 0.22%, Z = −3.107; *p* < 0.005; 16 Hz: 0.53 ± 0.12%, Z = −3.296; *p* < 0.005; 32 Hz: 0.29 ± 0.06%, Z = −3.296; *p* < 0.005; *n* = 14 from 9 animals) compared to the lowest frequency tested (0.5 Hz: 1.34 ± 0.30%). (**h**) Values for 10-90% rise time of the fluorescence responses of GACh2.0 expressing MEC stellate neurons to electric stimulations consisting of a train of 20 pulses at higher frequency (1 Hz: 8.1 ± 1.5 s, Z = 1.859; *p* = 0.06; 2 Hz: 6.2 ± 1.1 s, Z = 0.001; *p* = 0.99; 4 Hz: 3.8 ± 0.3 s; Z = −2.197; *p* < 0.05; 8 Hz: 2.4 ± 0.3 s, *Z* = −2.366; *p* < 0.05; 16 Hz: 2.1 ± 0.3 s, *Z* = −2.366; *p* < 0.05; *n* = 7 from 5 animals) compared to the lowest frequency tested (0.5 Hz: 6.5 ± 0.9 s), and values for decay time constant of the fluorescence responses of GACh2.0 expressing MEC stellate neurons to electric stimulations consisting of a train of 20 pulses at higher frequency (1 Hz: 30.4 ± 3.1 s, *Z* = 0.169; *p* = 0.87; 2 Hz: 30.8 ± 2.0 s, *Z* = 0.338; *p* = 0.74; 4 Hz: 33.0 ± 2.1 s; Z = 0.338; *p* = 0.74; 8 Hz: 36.4 ± 6.1 s, Z = 1.363; *p* = 0.17; 16 Hz: 31.0 ± 2.4 s, *Z* = 0.169; *p* = 0.87; *n* = 7 from 5 animals) compared to the lowest frequency tested (0.5 Hz: 30.8 ± 5.7 s). Note the stimulation frequency-dependent shortening in 10-90% rise time, but not decay time constant. (**i**) Fluorescence responses of GACh2.0 expressing MEC stellate neuron to electric stimulations consisting of a train of up to 80 pulses at 2 Hz. (**j**) Values for the maximal responses of GACh2.0 expressing MEC stellate neurons to electric stimulations consisting of a train of up to 80 pulses at 2 Hz (2 pulses: 0.70 ± 0.15%, *Z* = 1.960; *p* = 0.05; 5 pulses: 1.53 ± 0.38%, Z = 2.521; p<0.05; 10 pulses: 2.22 ± 0.56%, Z = 2.521; *p* < 0.05; 20 pulses: 3.29 ± 0.95%, Z = 2.521; *p* < 0.05; 40 pulses: 4.07 ± 1.22%, Z = 2.366; *p* < 0.05; 80 pulse: 3.65 ± 1.30%, *Z* = 2.366; *p* < 0.05; *n* = 8 from 4 animals) compared to single pulses (1 pulse: 0.50 ± 0.09%). (**k**) Values for 10-90% rise time of the maximal responses of GACh2.0 expressing MEC stellate neurons to electric stimulations consisting of a train of up to 80 pulses at 2 Hz (2 pulses: 1.8 ± 0.4 s, *Z* = 1.718; *p* = 0.09; 5 pulses: 2.0 ± 0.3 s, *Z =* 1.955; *p =* 0.05; 10 pulses: 3.3 ± 0.3 s, *Z =* 2.666; *p* < 0.01; 20 pulses: 5.0 ± 0.6 s, *Z* = 2.666; *p* < 0.01; 40 pulses: 6.5 ± 1.4 s, *Z* = 2.666; *p* < 0.01; *n* = 9 from 6 animals) compared to single pulses (1 pulse: 1.3 ± 0.3 s), and decay time constant of the maximal responses of GACh2.0 expressing MEC stellate neurons to electric stimulations consisting of a train of up to 80 pulses at 2 Hz (2 pulses: 32.2 ± 2.6 s, *Z* = −0.296; *p* = 0.77; 5 pulses: 33.9 ± 2.1 s, *Z* = 0.178; *p* = 0.86; 10 pulses: 32.8 ± 1.1 s, *Z* = −0.415; *p* = 0.68; 20 pulses: 32.7 ± 1.7 s, *Z* = −0.338; *p =* 0.75; 40 pulses: 31.8 ± 2.3 s, *Z* = −0.415; *p* = 0.68; *n* = 9 from 6 animals) compared to single pulses (1 pulse: 32.9 ± 3.5 s). Note the stimulation pulse number-dependent increase in 10-90% rise time, but not decay time constant. Asterisks indicate *p* < 0.05 (Wilcoxon tests).

To verify the cholinergic nature of electrically evoked ΔF/F responses, we included atropine in the bath solution. Unexpectedly, bath application of 1 μM atropine, which was sufficient to block ACh puff-evoked ΔF/F responses (**Fig. S9**), failed to block L1 stimulation-evoked ΔF/F responses, and at times, slightly enhanced ΔF/F responses in GACh2.0 expressing neurons (**Fig. S12b**). Although the atropine-induced enhancement of ΔF/F responses is surprising, the finding is consistent with previous reports of the atropine-potentiated ACh release attributed to a predominant presynaptic muscarinic autoreceptor inhibition mechanism^30–32^. Since presynaptic muscarinic receptors are primarily M2Rs and/or M4Rs^21, 33, 34^, we examined whether a similar presynaptic inhibition mechanism may exist in cholinergic transmission in MEC with (5*R*,6*R*)6-(3-propylthio-1,2,5-thiadiazol-4-yl)-1-azabicyclooctane (PTAC), which is both an agonist of M_2,4_Rs and an antagonist of M_1,3,5_Rs^35^. Bath application of 20 μM PTAC, which is sufficient to activate M_2,4_Rs and inhibit M_3_R-like GACh2.0, blocked the electrically evoked ΔF/F responses in GACh2.0 expressing stellate neurons (**Fig. S12c**). Together, these pharmacological findings are consistent with the view that a presynaptic M_2,4_R-mediated inhibition mechanism regulates ACh release in MEC.

To determine whether GACh sensors can report other modes of endogenous ACh release, we studied glutamatergic neurons in the medial habenula (**MHb**), which release ACh as a co-neurotransmitter during high frequency firing^8, 36^. To image the regulation of habenula cholinergic transmission, we made AAV viral expression of GACh2.0 in the interpeduncular nucleus (**IPN**), the major target of MHb cholinergic neurons. Two weeks after GACh2.0 expression, we prepared acute brain slices containing the source nucleus of MHb, the target IPN and the projections between the two areas carried by the fasciculus retroflexus (**MHb-fr-IPN**) (**Fig. S13a,b**). Two-photon imaging showed that brief 1-, 10-, 20- or 50-Hz 5-second electric stimuli evoked little change in fluorescence, whereas application of 100-Hz stimuli elicited small ΔF/F responses in GACh2.0 expressing IPN neurons (**Fig. S13e,g**). We have recently reported that GABA potentiates the optically evoked habenula cholinergic currents in IPN neurons via a presynaptic GABA_b_R-mediated heteroreceptor enhancement mechanism^29^. Consistent with those findings, GABA and baclofen, a GABA_b_ receptor agonist, enhanced ΔF/F responses in GACh2.0 expressing neurons by 6 fold, whereas saclofen, a GABA_b_ receptor antagonist, reversed the potentiative effects. Moreover, tiotropium completely abolished the potentiated ΔF/F responses, while donepezil, an acetylcholinesterase inhibitor^37^, prolonged the potentiated ΔF/F responses in GACh2.0 expressing neurons (**Fig. S13d, f, h-j**). These results confirm the presynaptic heteroreceptor enhancement mechanism in habenula cholinergic transmission with a GACh2.0-dependent optical approach. We next examined the effects of GABA on frequency-dependent ACh release in IPN. In the presence of 2 μM baclofen, 10-Hz electric stimuli evoked small ΔF/F responses in GACh2.0 expressing IPN neurons, which were progressively enhanced with increasing frequency from 20–100 Hz. These findings support the notion that in the presence of GABA in IPN, habenula neurons, which fire action potentials at frequencies up to 10–25 Hz *in vivo*^38^, can enhance cholinergic transmission to interpeduncular neurons, thought to be critical for fear control36. Together, the data collected from MEC and MHb-fr-IPN indicate that different firing patterns can create distinct ACh release modes in the central cholinergic system.

In addition to the control of ACh release, another unresolved fundamental question about cholinergic signaling is the extent of volume transmission. During the investigation of firing pattern-controlled ACh release, we noted that at times, the electrically evoked fluorescence responses exhibited obvious spatial heterogeneity across subcellular areas of GACh2.0 expressing L2 stellate neurons. Analysis of the minimal electric stimulation-evoked subcellular fluorescence responses in GACh2.0 expressing neurons revealed one or a few hot spots with largest ΔF/F responses, whereas other areas had smaller or undetectable changes in ΔF/F (Fig. 5), suggesting spatially restricted release and clearance of ACh. Plotting the size of ΔF/F responses against the distance from the spots with maximal ΔF/F responses yielded an estimated volume transmission spread length constant of 9.0 μm for cholinergic transmission (Fig. 5e).

**Figure 5:**
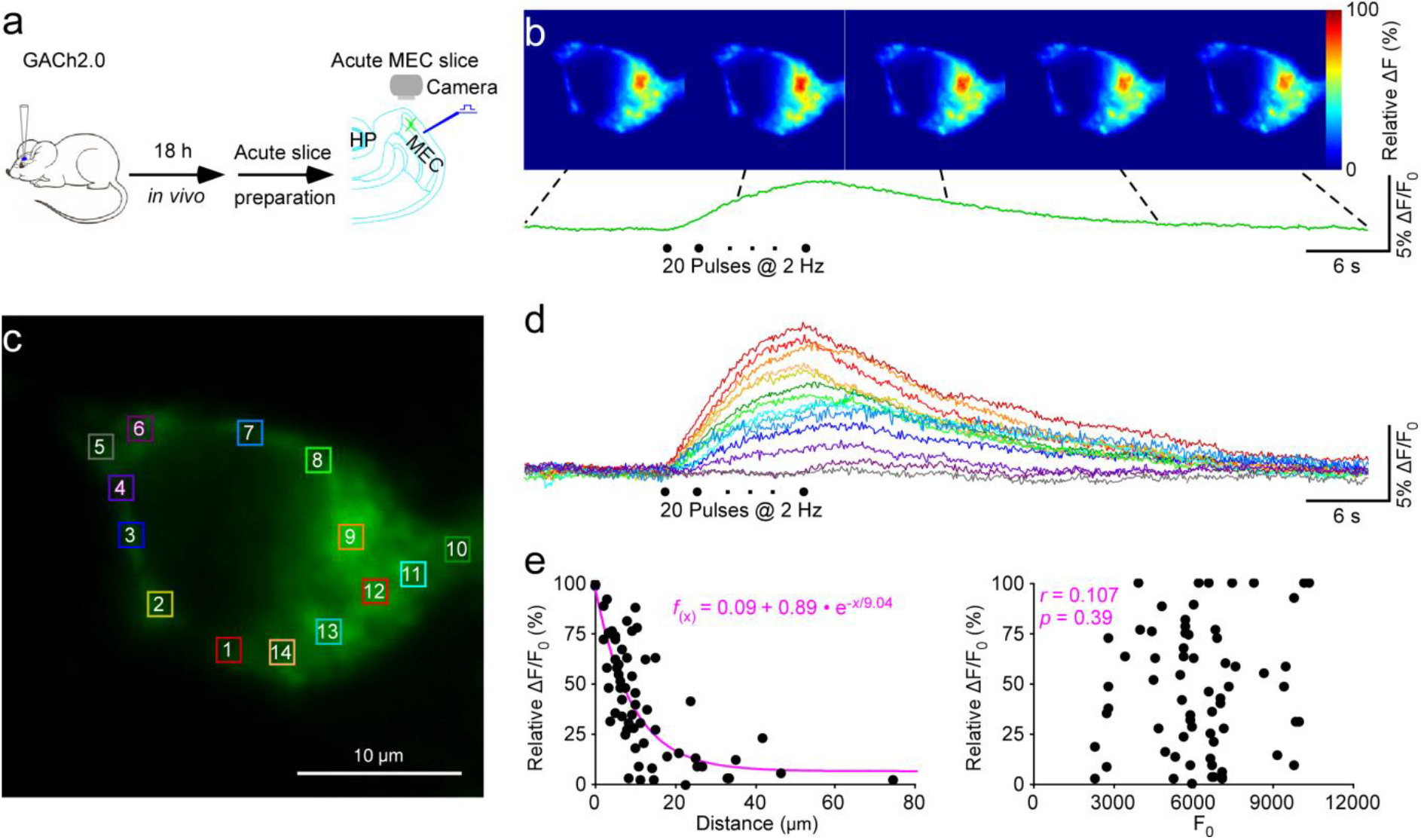
GACh2.0 illustrates subcellular cholinergic signals in MEC. (**a**) Schematic drawing outlines the design of stimulation-imaging experiments using an *in vivo* viral expression and *ex vivo* acute mouse MEC slice preparation. The inserted fluorescent image shows a GACh2.0 expressing MEC L2 stellate neuron. HP: hippocampus; MEC: medial entorhinal cortex. (**b**) Snapshots of relative fluorescence responses of the GACh2.0 expressing neuron to a minimal L1 electric stimulation. The lower fluorescence recording trace shows the average response of the neuron. (**c-d**) L1 electric stimulation-evoked fluorescence responses in local areas marked by color squares (~1.5 μm × ~1.5 μm) in c were shown in d. Note the near maximal responses seen in multiple areas suggestive of possible activation of multiple cholinergic fibers and/or release sites, and the slower rising times of smaller responses expected for diffused ACh. (**e**) Left, plot of the size of local ΔF/F responses against the distance from the area #1, which had the largest ΔF/F. The data points (n = 67 from 6 neurons of 6 animals) were arbitrarily fitted to a single exponential decay function (pink line), resulting in an estimated volume spread length constant of ~9 μm. Right, plot of the size of ΔF/F against the absolute F_0_ indicates no correlation between ΔF/F and F_0_ (*n* = 67; Normality test *p* = 0.06; Constant variance test *p* = 0.80; *r* = 0.107; *p* = 0.39; Linear regression *t* test).

Because the estimated cholinergic volume transmission size in MEC is rather small, we sought to independently verify our finding. Using the same *in vivo* Sindbis expression and *ex vivo* acute mouse slice preparation, we examined the minimal electric stimulation-evoked local fluorescence responses along the somatodendritic axis of GACh2.0 expressing CA1 neurons. Consistent with previous studies^25^, we found that electric stimuli of the stratum oriens and stratum pyramidale were most likely to elicit cholinergic responses in GACh2.0 expressing CA1 neurons. The largest fluorescence responses were typically observed at one or at most a few hot spots in somata of GACh2.0 expressing neurons, whereas fluorescence responses at other somatic and dendritic areas of the same neurons were much smaller or undetectable (**Fig. S14**). Plotting the size of ΔF/F responses against the distance from the maximal ΔF/F responses gave an estimated volume transmission spread length constant of 15.6 μm for the cholinergic transmission in CA1 neurons (**Fig. S14d**), confirming that the central cholinergic volume transmission is spatially restricted.

To test whether GACh sensors may be employed together with other optical techniques, we examined the feasibility of an all-optical approach, namely optogenetic activation as well as optical report of cholinergic transmission simultaneously. We again used the *in vivo* viral expression and *ex vivo* acute mouse slice preparation to examine fluorescence responses in neurons of MEC, one of the brain areas that receives the densest cholinergic inputs^26, 27^. Specifically, in ChAT-Cre mice, we first injected AAV virus into BF of ChAT-Cre mice to express DIO-oChIEF-tdTomato and allowed three weeks for the optogenetic probe to be expressed and transported to the terminal fields within MEC^39^. We then injected Sindbis virus to express GACh2.0 in MEC layer 2 stellate neurons MEC and allowed 18 hours for the sensor to be expressed before preparing acute entorhinal cortical slices. During the experiments, we employed single-photon laser pulses to optogenetically stimulate oChIEF-expressing cholinergic fibers in MEC, and simultaneously used two-photon laser scanning, which is insufficient to activate oChIEF-expressing fibers^40^, to image fluorescence responses in GACh2.0 expressing stellate neurons in MEC (**Fig. S15a**). Twenty 5-ms 473-nm single-photon laser pulses (at 1 Hz) elicited consistent fluorescence responses in GACh2.0 expressing neurons, which were largely blocked by bath application of 20 nM PTAC (**Fig. S15b-c**). These results establish the feasibility of all-optical activation and measurement of ACh release.

### Applications of GACh sensors in non-neuronal tissues

Given the role of peripheral and non-neuronal cholinergic signals in diverse physiological actions in a large variety of non-neuronal cells^5–7^, we explored the utility of GACh2.0 ACh reporting in some non-neuronal cells and tissues. ACh released from parasympathetic nerve terminals acts on the pancreas to trigger insulin secretion in response to food uptake^41^. We made Sindbis viral expression of GACh2.0 in the mouse pancreas *in vivo* for ~18 hrs, and then imaged fluorescence responses of GACh2.0 expressing cells in acutely prepared pancreas tissue slices (Fig. 6a). Single electric shocks of local parasympathetic cholinergic fibers evoked evident fluorescence responses in GACh2.0 expressing pancreatic cells (Fig. 6b,c; **Movie S5**). Furthermore, increasing the number of stimulation pulses delivered at 2 Hz linearly increased the amplitude of ΔF/F responses (Fig. 6d,e). As with central cholinergic transmission in MEC, bath application of 1 μM atropine failed to block, whereas 20 μM PTAC blocked ΔF/F responses in GACh2.0 expressing pancreatic cells (Fig. 6f,g), consistent with the engagement of a possible presynaptic inhibition mechanism. In addition, we tested the applicability of GACh2.0 in the mouse adrenal gland using a similar *in vivo* viral expression and acute tissue slice imaging approach (**Fig. S16a**). Similar to pancreastic cells, single local electric stimuli applied to activate cholinergic fibers that innervate the adrenal gland^42^ evoked detectable ΔF/F responses in GACh2.0 expressing adrenal cells (**Fig. S16b**; **Movie S6**). Increasing the number of stimulation pulses increased the amplitude of ΔF/F responses that plateaued after 10 pulses (**Fig. S16c**). Again, 1 μM of atropine failed to block, while 20 μM PTAC blocked ΔF/F responses in GACh2.0 expressing adrenal cells (**Fig. S16d**), again in agreement with a possible presynaptic inhibition mechanism. Together, these results suggest that GACh2.0 can monitor ACh in non-neuronal cells.

**Figure 6:**
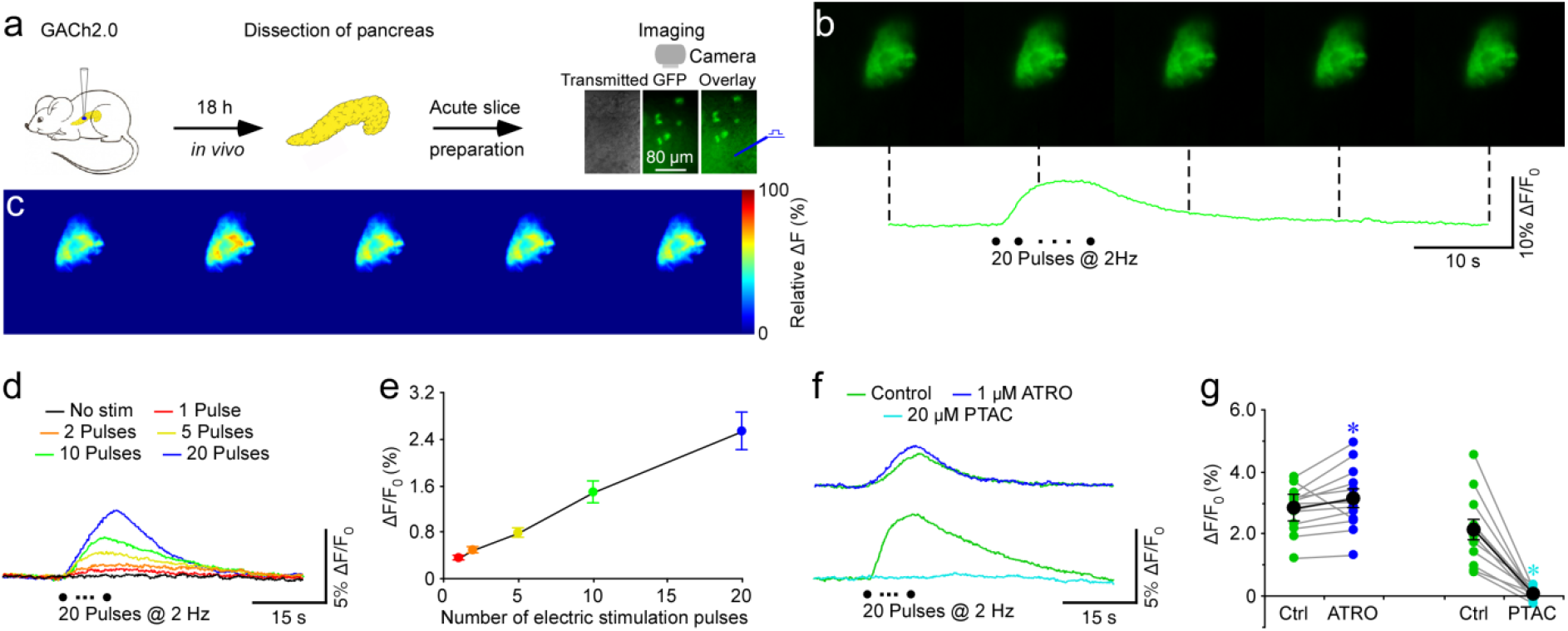
GACh2.0 permits visualization of cholinergic transmission in the pancreas. (**a**) Schematic drawing outlines the design of stimulation-imaging experiments using an *in vivo* viral expression and *in vitro* mouse pancreas tissue slice preparation. Inserts show transmitted light (left), fluorescence microscopic (middle) and overlay (right) images of GACh2.0 expressing pancreatic cells. (**b**) Snapshots of fluorescence responses of a GACh2.0 expressing pancreatic cell to local electric stimulation. (**c**) Relative fluorescence responses of the GACh2.0 expressing pancreatic cell to local electric stimulation shown in a heat map format. (**d**) Fluorescence responses of a GACh2.0 expressing pancreatic cell to electric stimulations consisting of a train of up to 20 pulses at 2 Hz. (**e**) Values for the maximal responses of GACh2.0 expressing pancreatic cells to electric stimulations consisting of a train of up to 20 pulses at 2 Hz (2 pulses: 0.49 ± 0.05%, *Z* = 3.148, *p* < 0.005; 5 pulses: 0.78±0.08%, Z = 3.527, *p* < 0.005; 10 pulses: 1.48 ± 0.19%, *Z* = 3.621, *p* < 0.005; 20 pulses: 2.53±0.22%, *Z* = 3.621, *p* < 0.005; *n* = 17 from 11 animals) compared to single pulses (1 pulse: 0.35±0.04%). (**f**) Electrically evoked fluorescence responses of GACh2.0 expressing pancreas cell bathed in the normal ACSF, in ACSF containing 1 μM atropine (ATRO) or 20 μM PTAC. (**g**) Values for the maximal evoked responses of GACh2.0 expressing pancreas cells in presence of ATRO (Ctrl: 2.79±0.20%; Exp: 3.13 ± 0.28%, *Z* = 2.631, *p* < 0.01; *n* = 13 from 8 animals) and PTAC (Ctrl: 2.11±0.33%; Exp: 0.05 ± 0.05%, *Z* = −3.059, *p* < 0.005; *n* = 12 from 5 animals) compared to control responses in normal ACSF. Black dots indicate average responses and asterisks indicate *p* < 0.05 (Wilcoxon tests).

### *Applications of GACh sensors* in transgenic *Drosophila in vivo*

Next, we tested whether GACh sensors could monitor functional cholinergic transmission in the intact brain of *Drosophila in vivo*. We performed two-photon imaging of odor-induced synaptic responses in the antennal lobe of *Drosophila*, which consists of powerful cholinergic synaptic inputs from olfactory receptor neurons to principal projection neurons^43^. We created UAS-GACh1.0 and -GACh2.0 transgenic flies, and then crossed them with a GH146-Gal4 driver line^44^ to selectively express GACh1.0 and GACh2.0 in antennal lobe projection neurons. In head-fixed living GACh transgenic flies, application of the odorant isoamyl acetate induced region-specific and dose-dependent ΔF/F responses in the DM2 glomerulus, that was not observed in the DA1 glomerulus (Fig. 7a-e; cf.^45, 46^). The control application of odor solvent, mineral oil alone did not evoke significant ΔF/F changes in GACh transgenic flies (Fig. 7b-c). As expected, the odor-evoked changes in ΔF/F in GACh2.0 transgenic flies were ~2-fold larger than those in GACh1.0 transgenic flies (Fig. 7e). Using the spectrum non-overlapping red Ca^2+^ indicator RGECO^47^, we re-examined the odor-induced effects in both control RGECO and GACh1.0/2.0::RGECO transgenic flies. We found the isoamyl acetate-induced Ca^2^+ transients in similar glomerular areas in control RGECO, GACh1.0::RGECO and GACh2.0::RGECO transgenic flies (**Fig. S17**), confirming the results obtained with GACh sensors. Further analysis showed that the isoamyl acetate-evoked Ca^2^+ transients in the glomerulus DM2 were indistinguishable between control RGECO, GACh1.0::RGECO and GACh2.0::RGECO transgenic flies (**Fig. S17**), suggesting that expression of GACh sensors had little effect on the endogenous receptor-mediated Gq-coupled Ca^2^+ signaling. Taken together, the results indicate that GACh can effectively monitor the dynamics of physiological ACh release in functional neuronal circuits in *Drosophila in vivo*.

**Figure 7:**
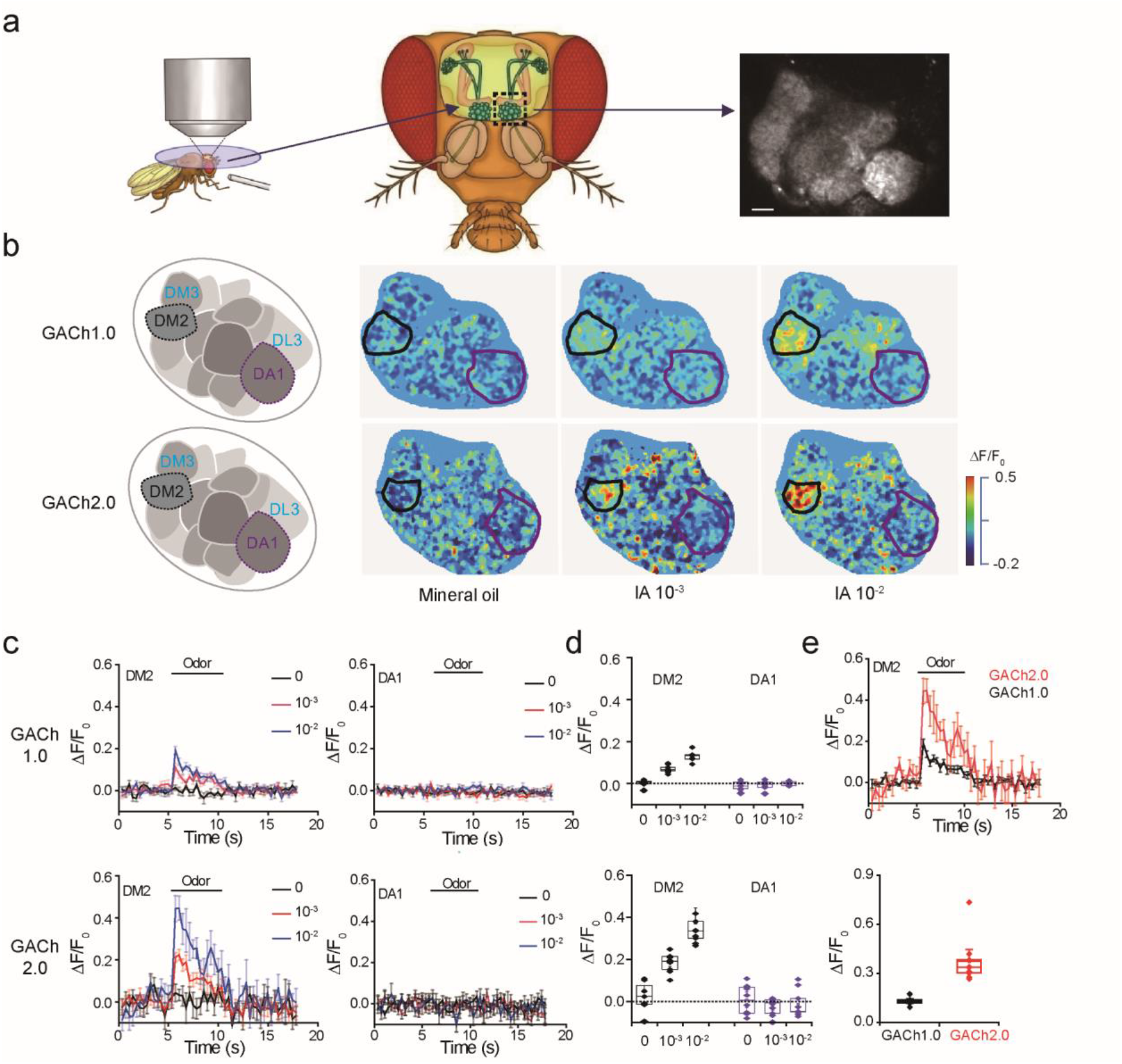
GACh sensors reveals dynamics of endogenous ACh release in *Drosophila*. (**a**) Schematic illustration of the two photon imaging setup of the *Drosophila* olfactory system. Odor was delivered near the antenna (Left) and GACh signals were measured in the antennal lobe area of GH146-Gal4: UAS-GACh flies (Right). Scale bar, 10 μm. (**b**) Pseudocolor images of GACh expressing antenna lobes show fluorescence responses to mineral oil and odor isoamyl acetate (IA). (**c**) Time courses of the lA-dependent responses in DM2 and DA1 glomeruli of GACh 1.0 (upper plots) and GACh2.0 (lower plots) expressing antenna lobes. (**d**) Values for the maximal ΔF/F_0_ in DM2 and DA1 glomeruli of GACh 1.0 (upper plot; DM2: 0.29 ± 0.49%, *n* = 9; DA1: −1.01 ± 0.68%, *n* = 9, *U* = 55, *p* = 0.22 for mineral oil; DM2: 7.02 ± 0.56%, *n* = 9; DA1: −0.61 ±0.80%, *n* = 9, *U* = 81, *p* < 0.001 for 10^−3^ IA; DM2: 12.97 ±1.28%, *n* = 7; DA1: 0.17 ± 0.39%, *n* = 7, *U* = 49, *p* < 0.005 for IA 10^−2^) and GACh2.0 (lower plot; DM2: 2.27 ± 2.34%, *n* = 8; DA1: −0.73 ± 1.41%, *n* = 8, *U* = 35, *p* = 0.80 for mineral oil, DM2: 18.78 ± 1.36%, *n* = 10; DA1: −2.37 ± 1.06%, *n* = 10, *U* = 100, *p* < 0.001 for 10^−3^ IA; DM2: 37.30 ±4.79%, *n* = 9; DA1: −0.55 ±2.68%, *n* = 9, *U* = 81, *p* < 0.001 for 10^−2^ IA) expressing antenna lobes (Mann-Whitney Rank Sum non-parametric test). (**e**) Upper, IA-evoked responses in DM2 glomerulus of GACh 1.0 and GACh2.0 transgenic flies. Lower: Values for the maximal ΔF/F_0_ in DM2 glomerulus of GACh 1.0 and GACh2.0 transgenic flies (GACh1.0: 12.97 ± 1.28%, *n* = 7; GACh2.0: 37.30 ± 4.79% *n* = 9; *U* = 63, *p* < 0.005, Mann-Whitney Rank Sum non-parametric test).

### Application of GACh sensors in mouse visual cortex in vivo

Finally, we tested whether GACh sensors were able to detect the endogenous ACh release in rodent brain *in vivo*. We used a visual attention model in a 2-photon setting of awake mice^48^. Specifically, GACh sensors were sparsely expressed by injection of Sindbis virus in visual cortex after behavioral habituation of mice, and *in vivo* two photon imaging experiments were conducted 16 h later with visual stimuli delivered by a monitor in front of the mice (Fig. 8a; **see methods**; cf.^49^). To monitor potential ACh release in the visual cortex, white filled circles expanding on the screen (10 s) were given as stimuli designed to engage visual attention, followed by 50 s darkness (Fig. 8b). Interestingly, we observed a subset of GACh2.0 expression neurons (ROI#1) repeatedly responded to the expanding circle with increase of fluorescence signals, consistent with ACh release upon attentional states^50, 51^, while other subset of neurons (ROI#2) remained quiescent in fluorescence (Fig. 8b,c), which may reflect a functional heterogeneity and complex cholinergic projection in the visual cortex. Analysis of multiple responding cells in different animals revealed a time-lock and statistically significant fluorescence response to the visual stimulation (Fig. 8d). Taken together, the GACh sensors were sensitive enough to report the attentional like visual stimuli evoked ACh release in awake behaving mice, which opens the door to study ACh function in many other brain regions in rodents *in vivo*.

**Figure 8:**
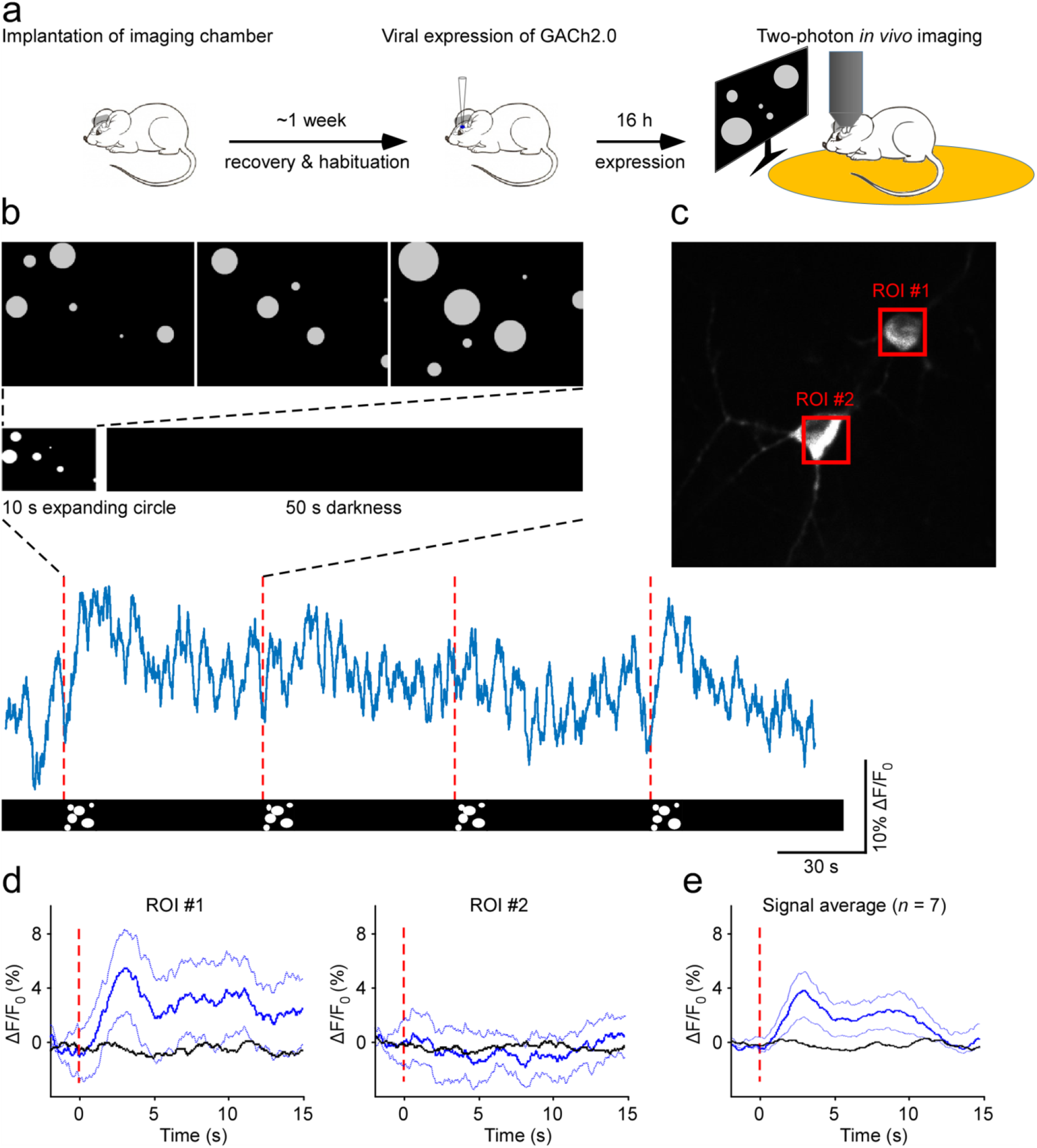
Attentional visual stimuli evoked ACh release in behaving mice. (**a**) Schematic drawing outlines the design of *in vivo* imaging experiments. (**b**) Upper, schematic representation of the visual stimulus to head-fixed behaving mice. The stimulus consisted of 10 seconds of expanding white circles appearing at random locations on the screen, followed by 50 seconds of darkness. Lower, 4-minute fluorescence response traces corresponding to four repetitions of the stimulus. (**c**) An imaged region (100 × 100 μm, 120 μm deep) contains two GACh2.0 expressing neurons with the red squares indicating regions of interests (ROIs). (**d**) Mean fluorescence responses from ROIs shown in (b). The fluorescence response traces were divided into 10 second segments with every minute containing one 10-s trace corresponding to the period of visual stimulation and five 10-s traces corresponding to periods of darkness. The signal from the middle three dark segments (black line) was compared to the signal from the visual stimulus segments (blue line; 15-20 trials per region). Dotted lines around the solid blue trace show the 95% confidence interval obtained by bootstrap. Note that ROI #1, but not ROI #2, shows an increase in fluorescence responses to visual stimulus. (**e**) Average fluorescence responses obtained during the period of visual stimulation compared to those during the period of darkness (Visual: 3.65 ± 2.42%; Dark: 0.04 ± 0.65%; *Z* = −2.366, *p* < 0.05; *n* = 7 from 4 animals).

## DISCUSSION

In this study, we have developed and validated a family of genetically-encoded fluorescent ACh probes, or GACh sensors. GACh sensors have the sensitivity, ligand specificity, SNR, kinetics and photostability, the properties that are suitable for monitoring cholinergic signals in diverse tissue preparations *in vitro, ex vivo* and *in vivo*. Using the new GACh sensors, we revealed a few key features of cholinergic transmission, including firing pattern-dependent release and spatially restricted volume transmission. We also demonstrated the feasibility of utilizing GACh sensors to visualize endogenous ACh transmission in both neuronal and non-neuronal tissues, providing the possibility for deciphering cholinergic mechanisms in diverse physiological and pathological conditions.

The genetically-encoded GACh sensors represent a new toolbox suitable for a wide-range of preparations. We show here that the sensors can be expressed in a dozen distinct types of cells (i.e., HEK293T cells, cultured cortical neurons, hippocampal CA1 and CA3 pyramidal neurons, entorhinal cortical L1 interneurons and L2 stellate neurons, barrel cortical L5 pyramidal neurons, interpeduncular neurons, visual cortex, and pancreas and adrenal cells in the mouse, thalamocortical neurons and thalamic reticular neurons in the rat, and olfactory projection neurons in the fly) using transfection, viral expression (AAV, Sindbis and lenti-virus) or transgenic techniques. In all the preparations, GACh sensors effectively detected exogenous and/or endogenous ACh with high specificity, and the ACh-evoked optical signals could be captured with epi-fluorescent, confocal and/or two-photon microscopy *in vitro* and *in vivo*. Overall, GACh sensors have EC_50_ equivalent to ~1 μM, ΔF/F sensitivity of up to ~100%, activation/inactivation kinetics within sub-second, and photostability capability to repeatedly report ACh for hours. We note that GACh sensors have a weak coupling to the downstream G protein intracellular signaling in cultured cells, yet this persistent weak coupling has no detectable effect on basic membrane properties, synaptic properties and cholinergic transmission in rodent neurons *in vitro*, or the sensory input-evoked cholinergic responses in fly and attention induced ACh release in mice visual cortex *in vivo*. We expect additional site-directed mutagenesis and screening to generate successor GACh sensors with even better performance (e.g., improved sensitivity and further reduced intracellular signaling generation) in the near future.

With the newly developed GACh sensors, we have investigated a few fundamental questions of cholinergic transmission. Central cholinergic neurons exhibit multiple distinct action potential firing patterns^28, 29, 38^, yet the functional significance of these firing patterns remains elusive. Here we report that BF cholinergic neurons use low frequency 0.5-2 Hz tonic firing to generate large plateau-like postsynaptic ACh signals, and 8-12 Hz theta rhythmic phasic firing to elicit small transient postsynaptic ACh signals. High-frequency activation of cholinergic fibers seems to be more effective in recruiting presynaptic auto-receptor inhibition mechanism to suppress ACh release^32^. As such, the low frequency firing-induced releases may be less suppressed and thereby generate plateau-like ACh signals, whereas the high frequency firing-induced release seems to yield more transient ACh signals. In addition, habenula neurons can fire high frequency action potentials of up to 10-25 Hz^35^. The firing triggers co-release of ACh with its primary neurotransmitter glutamate when a presynaptic GABA_b_-R-mediated enhancement mechanism is engaged. Although the detailed aspects of this regulation remain to be further dissected, our data are consistent with the view that presynaptic regulatory mechanisms (i.e., autoreceptor inhibition and heteroreceptor enhancement) may play key roles in governing release modes in central cholinergic transmission.

In addition, a long-standing question of central cholinergic transmission concerns the extent of volume transmission; some investigators propose that ACh acts globally affecting a large number of neurons, whereas others consider cholinergic volume transmission to be be more restricted^1–3, 9^. The new ACh sensors have allowed us to directly visualize the spread of released ACh in the hippocampus and MEC, and estimate cholinergic volume transmission spread length constant of ~10-15 μm. Because the minimal electric stimulation can still activate multiple ACh release sites, the calculated volume transmission spread length constants may be overestimated. Nevertheless, these estimations provide the first suggestion that central cholinergic volume transmission may have single-cell or subcellular specificity. Together, fine firing frequency-controlled release and spatially restricted volume transmission advance our fundamental understanding of the control and precision of cholinergic signaling.

Our ability to visualize ACh signals in rodent models *ex vivo* and *in vivo* aided by the newly developed ACh sensors should help to advance our understanding of the pathogenesis of various diseases. For example, cholinergic signals are essential for high-level cognitive functions, including learning and memory, and dysregulation of cholinergic transmission is observed in many neurological disorders, including Alzheimer’s disease. However, the cholinergic hypothesis-based acetylcholinesterase inhibitor treatment, the only available therapy for Alzheimer’s disease^33^, has limited efficacy and thus is far from ideal^37, 52^. Further understanding the mechanisms of central cholinergic transmission in physiological and pathological conditions is central to development of effective therapeutic strategies for Alzheimer’s disease. Moreover, defective cholinergic signals have been implicated in the pathophysiology and treatment of a number of other non-neurological diseases^5–7^. For instance, ACh modulates insulin secretion from the pancreatic islets of Langerhans^41^ and the activity of M_3_Rs is essential for glucose tolerance and insulin release^34, 53^. Moreover, ACh controls release of hormones in the adrenal critical for regulation of stress and blood pressure^42^. Last but not least, cholinergic signals seem to control the progress of inflammation^54^ and tumorigenesis^55, 56^. We show here that GACh sensors are effective in monitoring cholinergic transmission in non-neuronal cells as well, including the pancreas and the adrenal, endorsing the use of this new toolbox to unravel the cholinergic mechanisms underlying these medical conditions.

## METHODS

### Animal preparation

Male and female Sprague Dawley rats, and wild type and ChAT-Cre transgenic C57BL/6 mice were used to prepare cultured neurons, cultured hippocampal slices, acute brain slices, acute pancreas and adrenal slices in this study. Animals were maintained in the animal facilities at the Peking University, the National Institute of Biological Sciences, Beijing, China, University of Southern California, Stony Brook University or the University of Virginia, and family or pair housed in the temperature-controlled animal room with 12-h/12-h light/dark cycle. All procedures for animal surgery and maintenance were performed following protocols approved by the Animal Care & Use Committee of the Peking University, the National Institute of Biological Sciences, Beijing, China, University of Southern California, Stony Brook University or the University of Virginia and in accordance with US National Institutes of Health guidelines.

### Preparations of cultured cells, cultured neurons and cultured slices

HEK293T were cultured in DMEM (Gibco, MA) with 5% FBS (North TZ-Biotech Develop Co., Ltd, Beijing, China) at 37 °C, 5% CO_2_, and passed to poly-D-lysine (Sigma-Aldrich, MO) coated 12-mm glass coverslips in 24 well plates. Rat cortical neurons were prepared from postnatal 1-day old (P1) Sprague-Dawley rats as previously described^45^. The rat brains were dissected and digested by 0.25% Trypsin-EDTA (Gibco), and placed on to poly-D-lysine coated coverslips with density of 0.5-1×10^6^ cells/ml.

Cultured slices were prepared from P6-7 rats or mice following our previous studies^57, 58^. In brief, the hippocampi were dissected out in ice-cold HEPES-buffered Hanks’ solution (pH 7.35) under sterile conditions, sectioned into 400 μm slices on a tissue chopper, and explanted onto a Millicell-CM membrane (0.4-μm pore size; Millipore, MA). The membranes were then placed in 750 μl of MEM culture medium, contained (in mM): HEPES 30, heat-inactivated horse serum 20%, glutamine 1.4, D-glucose 16.25, NaHCO_3_ 5, CaCl_2_ 1, MgSO_4_ 2, insulin 1 mg/ml, ascorbic acid 0.012% at pH 7.28 and osmolarity 320. Cultured slices were maintained at 35°C, in a humidified incubator (ambient air enriched with 5% CO_2_).

### Preparations of acute tissue slices

Acute thalamic, barrel cortical, entorhinal cortical, hippocampal and MHb-fr-IPN brain slices, pancreas and adrenal tissues slices were prepared from P25-60 animals deeply anesthetized by xylazine-ketamine or pentobarbital (100 mg/kg) as described in our previous reports^36, 58^. The animals were decapitated and the brain block containing the thalamus, barrel cortex, MEC and/or hippocampus, the pancreas, or the adrenal was quickly removed and placed into cold (0-4°C) oxygenated physiological solution containing (in mM): 125 NaCl, 2.5 KCl, 1.25 NaH_2_PO_4_, 25 NaHCO_3_, 1 MgCl_2_, 25 dextrose, and 2 CaCl_2_, pH 7.4. The brain blocks were directly sectioned into 400-μm-thick brain slices using a DSK microslicer (Ted Pella Inc.) or a VT1200 vibratome (Leica, Germany), while the pancreas and adrenal were first embedded in low–melting temperature agar (2.5% in BBS) and then sectioned into 400-μm-thick tissue slices^59^. The tissue slices were kept at 37.0±0.5°C in oxygenated physiological solution for ~0.5-1 hr before imaging. During the recording and/or imaging the slices were submerged in a chamber and stabilized with a fine nylon net attached to a platinum ring. The recording chamber was perfused with oxygenated physiological solution. The half-time for the bath solution exchange was ~6 s, and the temperature of the bath solution was maintained at 34.0 ± 0.5 °C. All antagonists were bath applied.

### Molecular biology

Molecular cloning was typically carried out using the Gibson assembly^60^ with ~30-overlapping base primers and the Phusion DNA polymerase (New England Biolabs, MA), and verified by Sanger sequencing using in an in-house facility (sequencing platform in the School of Life Sciences of the Peking University). The chimeric GACh constructs were generated by subcloning full-length human GPCR cDNAs (hORFeome database 8.1, the Dana-Farber Cancer Institute Center for Cancer Systems Biology) into the pDisplay vector (Invitrogen, MA), with an IgK leader sequence inserted before the coding region and a stop codon inserted before the PDGFR transmembrane domain. The ICL_3_ of M1-5Rs, defined according to the UNIPROT database (**Fig. S1**), was replaced by a shorter cpGFP-inserted ICL_3_ from a β2 adrenergic receptor. The site-directed mutagenesis of the sequences of the two- and five-amino acid linkers in the N and C termini of cpGFP was made using primers containing various lengths of tr-inucleotides NNB (20 possible amino acids, Sangon Biotech, Shanghai, China). The applicable GACh sensors were then subcloned into the Sindbis viral vector, the lentiviral vector or the AAV package vector under the human synapsin promoter to ensure the neuronal expression. To create the transgenic *Drosophila*, fragments of GACh sensors including the IgK leader sequence were cloned into the pUAST vector, and subsequently injected into *Drosophila* embryo following a standard protocol (Fungene biotechnology, Beijing). To report the receptor endocytosis, super ecliptic pHluorin^43^ was cloned to the N terminus of M_3_R, with a three amino acid linker (GGA) to ensure correct protein folding and trafficking.

### Expression of GACh sensors and other recombinant proteins

HEK293T cells were typically transfected using the PEI method (with a typical ratio of 1 μg DNA:4 μg PEI), media replaced 4-6 h later, and imaged 24 h later. Cultured neurons were transfected after 7-9 days *in vitro* using the calcium phosphate transfection method and experiments were performed 48 h after transfection. Neurons in hippocampal cultured slices were infected after 8–18 days *in vitro* with lentivirus or Sindbis virus, and then incubated on culture media and 5% CO_2_ before experiments. For *in vivo* expression, P28-84 animals were initially anesthetized by an intraperitoneal injection of 2,2,2-Tribromoethanol (Avetin, 500 mg/kg) or ketamine and xylazine (10 and 2 mg/kg, respectively), and then placed in a stereotaxic frame. In some of the animals, AAV of GACh sensors (with a titer of >10^12^/ml) was injected into IPN with a microsyringe pump (Nanoliter 2000 Injector, WPI) using the previously described coordinates (AP: −3.13 mm from Bregma; DV: −4.95 mm; ML: 1.33 mm with 15° angle towards the midline)8, 36. Two weeks after the viral injection, acute MHb-fr-IPN brain slices were prepared from these for experiments. In other animals, a glass pipette was used to penetrate into the thalamic ventrobasal nucleus, thalamic reticular nucleus, the barrel cortex and MEC according to stereotaxic coordinates, or the dissected pancreas and adrenal, to deliver ~50 nl of viral solution by pressure injection to infect neurons, or pancreas and adrenal cells with GACh sensors. Acute brain or tissue slices of 400 μm thick were then prepared from these animals ~18 h later to carry out experiments. In the rest animals, or ChAT-Cre transgenic mice, AAV of DIO-oChIEF-tdTomato was first injected into BR according to the previously described coordinates^39^, and three weeks later, Sindbis virus of GACh sensors was injected into the MEC for ~18 h before preparing acute brain slices for experiments.

### Electrophysiology

Simultaneous dual whole-cell recordings were obtained from two nearby infected and non-infected hippocampal CA1 pyramidal neurons under visual guidance using fluorescence and transmitted light illumination^40, 58^. The patch recording pipettes (4-7 MΩ) were filled with intracellular solution containing 115 mM cesium methanesulfonate, 20 mM CsCl, 10 mM HEPES, 2.5 mM MgCl_2_, 4 mM Na_2_ATP, 0.4 mM Na_3_GTP, 10 mM sodium phosphocreatine, 0.6 mM EGTA, and 0.1 mM spermine and 0.5% biocytin (pH 7.25) for voltage-clamp recordings, or containing 120 mM potassium gluconate, 4 mM KCl, 10 mM HEPES, 4 mM MgATP, 0.3 mM Na_3_GTP, 10 mM sodium phosphocreatine and 0.5% biocytin (pH 7.25) for current-clamp recordings. Bath solution (29±1.5°C) contained (in mM): NaCl 119, KCl 2.5, CaCl_2_ 4, MgCl_2_ 4, NaHCO_3_ 26, NaH_2_PO_4_ 1, glucose 11, picrotoxin (PTX) 0.1, bicuculline 0.01, and 2-chloroadenosine 0.002, at pH 7.4 and gassed with 5% CO_2_/95% O_2_. PTX was excluded when GABA responses were examined. Whole-cell recordings were made with up to two Axoclamp 2B or Axopatch-200B patch clamp amplifiers (Molecular Devices, Sunnyvale, CA). Junction potentials were not corrected. Synaptic responses were evoked by bipolar electrodes with single voltage pulses (200 μs, up to 20 V). Synaptic AMPA and NMDA responses at −60 mV and +40 mV or GABA responses at 0 mV were averaged over 90 trials. To minimize the effect from AMPA responses, the peak NMDA responses at +40 mV were measured after digital subtraction of estimated AMPA responses at +40 mV. Cholinergic fibers in tissue slices were stimulated with a bipolar electrode placed ~50–200 μm from imaged cells with single or a train of voltage pulses (500 μs, up to 50 V) to evoke ACh release. Optogenetic activation of oChIEF-expressing cholinergic fibers was made with a train of 5-ms light pulses delivered at 1 Hz from a 470-nm blue M470F1 LED (Thorlabs, NJ). The blue light of the LED was fiber-coupled to an Ø200 μm fiber optic cannula positioned ~250 μm away from imaged neurons. The light power out of the cannula was set at 2 mW.

### Fluorescence imaging of cultured cells and neurons

HEK293T cells transfected with the muscarinic receptor-based chimeric constructs were imaged with a TECAN Safire2 fluorescence plate reader (TECAN, Männedorf, Switzerland; excitation, 480 nm; emission, 520 nm). During the measurement, the culture media was replaced with 100 μl Tyrode solution containing ACh at varied concentrations from 0-100 μM. The ΔF/F of each construct was obtained by averaging the ACh-induced fluorescence responses of 2-6 transfected wells (>100 cells/well) after digitally subtracting that of neighboring control non-transfected wells.

HEK293T cells and cultured neurons were perfused with standard extracellular Tyrode solution containing (in mM): 150 NaCl, 4 KCl, 2 MgCl_2_, 2 CaCl_2_, 10 HEPES and 10 Glucose, with pH of 7.4, in an imaging chamber during imaging. Agonist acetylcholine (Solarbio, Beijing, China), tiotropium bromide (Dexinjia Bio & Tech Co., Ltd, Jinan, China), isoamyl acetate and AF-DX 384 (Sigma-Aldrich) were delivered with a custom-made perfusion system and/or bath applied. The chamber was washed with Tyrode solution between applications and cleaned with 75% ethanol between experiments. HEK293T cells were imaged using an IX81 inverted microscope (Olympus, Tokyo, Japan) with a 40x/1.35 NA oil objective, a 475/28 excitation filter for cpGFP excitation, a 506LP dichroic mirror and a 515LP emission filter for imaging collection. The data were acquired with a Zyla sCMOS DG-152V-C1e-FI camera (Andor Technology, UK) controlled by μManager. Cultured neurons were imaged using an inverted Nikon Ti-E A1 confocal microscope (Nikon, Tokyo, Japan) with a 40x/1.35 NA oil objective and a 488 nm laser illumination.

### Fluorescence imaging of cells in cultured and acute slice preparations

Wide field epifluorescence imaging was performed using Hamamatsu ORCA FLASH4.0 camera (Hamamatsu Photonics, Japan) and GACh expressing cells in cultured hippocampal slices and acutely prepared brain slices are excited by a 460-nm ultrahigh-power low-noise LED (Prizmatix, Givat-Shmuel, Israel). The frame rate of FLASH4.0 camera was set to 10 Hz. To synchronize image capture with drug perfusion, electrical stimulation, and/or electrophysiological recording, the camera was set to external trigger mode and triggered by a custom-written IGOR Pro 6 program (WaveMetrics, Lake Oswego, OR). Agonists or antagonists, including acetylcholine and atropine (Sigma-Aldrich), and nicotine, oxotremorine M, TAPC and TMPH (Tocris Bioscience, Bristol, UK), were either bath applied or puff applied with a glass pipette positioned above the imaged neurons by 500-ms pressure pulses.

Two-photon imaging was performed using a custom-built microscope or an Olympus FV1000 microscope (for IPN experiments; Olympus, Japan). The parameters of frame scan were typically set at a size of 200 × 200 pixels and a speed of 1-2.5 frame/s. The fluorescence of oChIEF-tdTomato and GACh2.0 was excited by a femtosecond Ti:Sapphire laser (Chameleon Ultra II, Coherent) at a wavelength of 950 nm. Changes in fluorescence were quantified as increases in fluorescence from baseline divided by resting fluorescence (ΔF/F_0_) and averaged for ~10 trials. To quantify surface expression of GACh sensors, lentiviral expression of GACh1.0, GACh1.5 or GACh2.0 was made in the CA1 region of organotypic hippocampal cultured slices. About ~1-2 weeks after expression, GACh expressing CA1 pyramidal neurons were patch-clamp recorded and loaded with 5 μM Alexa Fluor 594 (Life Technologies) for ~10 minutes, and two-photon images were then taken at different compartments along the apical dendrites. The multiple patch-clamp recordings, optogenetics, epifluorescence and two-photon imaging were typically operated by a single custom-written IGOR Pro 6 program (WaveMetrics, Lake Oswego, OR)40.

### Immunocytochemistry

Mice infected with GACh sensors were deeply anesthetized with pentobarbital (400 mg/kg; i.p.), and transcardially perfused first with cold normal saline and then 4% paraformaldehyde in 0.1 M PBS. Brain blocks were post-fixed for ⩾4 hr, cryoprotected in 30% sucrose for ⩾24 h, then embedded in tissue freezing medium and sectioned into 50-μm-thick coronal sections with a freezing Leica CM 1900 microtome (Leica, Germany). To label cholinergic terminals from MHb and GACh expressing neurons in IPN, tissue sections were rinsed and immunoreacted with Goat ChAT antibody (1:500, Millipore, ab144p) and Rabbit GFP antibody (1:500, Abcam, #ab6556), and then labeled with goat-anti-rabbit second antibody conjugated Alexa 488 and donkey-anti-goat second antibody conjugated Alexa 555 after extensive washing. The immunolabeled tissue sections were imaged with a confocal microscope.

To recover the morphology of recorded neurons, the slices were fixed by immersion in 3% acrolein/4% paraformaldehyde in 0.1 M PBS at 4°C for 24 hrs after *in vitro* patch-clamp recordings with internal solution containing additional 1% biocytin, and then processed with the avidin-biotin-peroxidase method to reveal cell morphology. The morphologically recovered cells were examined and reconstructed with the aid of a microscope equipped with a computerized reconstruction system Neurolucida (MicroBrightField, Colchester, VT).

### Fluorescence imaging of transgenic Drosophila

Transgenic *Drosophila* lines with strong GACh expression levels and robust odor responses were chosen after crossing UAS-GACh1.0 and -GACh2.0 transgenic flies with a GH146-Gal4 driver line^44^. They were reared at room temperature for 8~12 days on standard medium after eclosion before experiments. For imaging experiments, the mounting of the live flies was essentially similar as described before^61^. Live flies were mounted to a small dish, with their rectangular patch of cuticle between the eyes, excessive fat bodies and air sacs surrounding the antennal lobe removed, and the pair of muscles underneath the proboscis cut to reduce the brain movement. The exposed brains were perfused (at 0.5 ml/min) with adult-like hemolymph (ALH) containing (in mM): 108 NaCl, 5 KCl, 5 HEPES, 5 Trehalose, 5 sucrose, 26 NaHCO_3_, 1 NaH_2_PO_4_, 2 CaCl_2_ and 1~2 MgCl_2_, with osmolarity 275 mOsm. 5% CO_2_/ 95% O_2_ were continuously bubbled to maintain ALH at pH 7.3. Isoamyl acetate (Sigma-Aldrich; Cat# 306967) was initially diluted by 100-fold or 1000-fold (vol/vol) in mineral oil (Sigma-Aldrich; Cat# 69794) and then placed in a glass bottle (100 μ l in 900 μl mineral oil), delivered at 200 ml/min, and mixed with purified air (1000 ml/min). The mixed air stream was presented to flies through a 1-cm-wide opening Teflon tube placed ~1 cm from their antennas, and controlled by a Teflon solenoid valves and synchronized with the image acquisition system by Arduino boards. Imaging was made using a commercial Olympus BX61WI two-photon microscope with a 25x NA: 1.05 water-immersion objective and a mode-locked Ti:Sapphirelaser (Mai tai) tuned to 950 nm. The Glomeruli were identified according to the previous established antennal lobe map^50^.

### Surgery for in vivo imaging in awake mouse

The surgery was performed in 2 steps. First, we removed the skin on top of the head and attached the metal recording chamber. The mouse was then given 3-5 days to recover, after which it was habituated to head-fixation. Approximately 1 week after the initial surgery, we opened the skull over the primary visual cortex (centered ~2.5 lateral, ~1.5 mm anterior from lambda), and did several 100 nl pressure injections of 1/5 diluted virus in the posterior half of the opening. The craniotomy was fitted by a cranial window made from 3 mm circular coverslip and a 2 × 2 mm square cut out of a #2 coverslip, as described previously^49^.

### Two-photon imaging and analysis

Imaging was performed starting 14-16 hours after the surgery. The awake mouse was head-fixed on circular treadmill and imaged using a custom built 2-p system using an InSight DS+ laser (Spectra Physics) and the ScanImage 5.1 software^62^. Images were acquired from individual cells (or small groups of cells when possible) continuously at either 30 Hz (512 × 512 pixels) or 60 Hz (256 × 256 pixels). The mouse was shown a stimulus consisting of 50 seconds of darkness followed by 10 seconds of expanding white circles appearing at random positions on the screen. All data analysis was done in Matlab (Mathworks). After automatic image alignment, each recording was manually inspected to make sure alignment is artifact free. ROIs were then manually drawn over the cell bodies and raw fluorescent traces were extracted. Fluorescent traces were filtered by a 2 second moving average window to reduce fluctuations. The fluoresce traces were then divided into 10 second segments corresponding to either periods of darkness or periods of visual stimulation and dF/F_0_ was calculated for each of them. The maximum dF/F_0_ was compared for periods with or without stimulation using a Wilcoxon rank sum test.

### Statistical analysis

Statistical results were reported as mean±s.e.m. Animals or cells were randomly assigned into control or experimental groups and investigators were blinded to experiment treatments. Given the negative correlation between the variation and square root of sample number, *n*, the group sample size was typically set to be ~10-25 to optimize the power of statistical tests and efficiency. Statistical significances of the means (*p*<0.05; two sides) were determined using Wilcoxon and Mann-Whitney Rank Sum non-parametric tests for paired and unpaired samples, respectively. Statistical significances of the linear relationships of two data groups were determined using linear regression *t* tests provided the normality and constant variance tests passed.

## AUTHOR CONTRIBUTIONS

J.J. Z. and Y.L. conceived the project. M.J. did GACh screening and optimization as well as its validation in cultured neurons and IPN slices. Y.L., Y.S. and H.J. designed and performed the work on transgenic flies. M.J. and J.F. preformed experiments related to calcium imaging, GPCR internalization Tango assay and FRET measurements. L. M. and L.Z. did *in vivo* imaging of GACh sensors in mouse visual cortex. M.L. supervised the imaging experiments on MHb-IPN brain slices. All authors contributed to data analysis. M.J., J.J. Z. and Y.L. wrote the manuscript with input from other authors.

## Acknowledgments

We thank Y. Rao for generous sharing of two-photon microscopy. We are also grateful to LQ Luo, S. Owen, Y. Rao and L. Nevin for critical reading of the manuscript. We thank Z. Ye for the help in art designing. This work was supported by the National Basic Research Program of China (973 Program; grant 2015CB856402) and The General Program of National Natural Science Foundation of China (project 31671118 and project 31371442) and the Junior Thousand Talents Program of China to Y.L.

